# Discovery of a Human Metabolite that Mimics the Bacterial Quorum-Sensing Autoinducer AI-2

**DOI:** 10.64898/2026.01.16.699917

**Authors:** Emilee E. Shine, Julie S. Valastyan, Vanessa Y. Ying, Jonathan Z. Huang, Mohammad R. Seyedsayamdost, Bonnie L. Bassler

## Abstract

Bacteria use small molecules to orchestrate collective behaviors in a process called quorum sensing (QS), which relies on the production, release, and group-wide detection of extracellular signal molecules referred to as autoinducers. One QS autoinducer, termed AI-2, is broadly used for inter-species bacterial communication, including in the mammalian gut. AI-2 consists of a family of interconverting compounds and adducts originating from 4,5-hydroxy-2,3-pentanedione. This complex speciation, coupled with the inherent instability of AI-2 congeners, have complicated isolation efforts. It has been known that mammalian epithelial cells produce an AI-2 mimic to which bacteria respond. However, the identity of the AI-2 mimic has remained elusive, presumably due to its instability, similar to that of known AI-2 compounds. Here, we developed a reactivity-based metabolomics approach to capture and identify a mammalian AI-2 mimic. Using a chemical strategy targeted at the α-diketone moiety of known AI-2s, we identify the unusual sugar L-xylosone, as well as the related metabolite L-xylulose, as AI-2 mimics. While L-xylulose is a common and naturally occurring sugar known in human metabolism, L-xylosone is a rare and highly reactive oxidation product. We established a facile synthetic route to access pure enantiomers of xylosone and confirmed that, like AI-2, the L-configuration is required for recognition by the bacterial AI-2 receptor, LuxP, whereas D-xylosone is inactive. L-xylosone is new to the human metabolome, suggesting that other chemically reactive small molecules that mediate host-microbe interactions await discovery. The identification of L-xylosone expands the AI-2 family of molecules and adds a new word to the lexicon of host-bacterial interactions.

## Introduction

The human gut microbiome consists of thousands of bacterial species that profoundly influence health and disease.^1–3^ Collectively, gut microbes possess far greater metabolic capacity than the host and synthesize a diverse array of metabolites, the identities of which continue to be uncovered. Some have been shown to drive microbe-microbe^4, 5^ and host-microbe interactions,^6, 7^ but the roles of many of these microbially-derived metabolites remain to be defined. The molecular mechanisms underlying these chemically-mediated interactions, which can be both beneficial or detrimental to human health, is another topic of intense interest. Microbiome metabolites are often challenging to identify in biological matrices, due to low abundance,^8, 9^ or chemical instability.^5, 10–12^ Advances in mass spectrometry have provided high-throughput routes to compound detection.^13–16^ Nonetheless, it remains challenging to assign structures to the majority of ions detected in metabolomics datasets.^17, 18^ Connecting function and regulation to known metabolites provides a further hurdle in unraveling chemically mediated interactions in the microbiome.

Small molecules underpin the cell-to-cell communication process called quorum sensing (QS), which enables bacteria to orchestrate collective behaviors. QS involves production, release, accumulation, and group-wide detection of extracellular signal molecules called autoinducers.^19, 20^ QS controls numerous collective behaviors in diverse bacteria, including bioluminescence, virulence factor production, antibiotic biosynthesis, and biofilm formation. There is increasing evidence that QS sensing is fundamental to interactions occurring in the context of the human microbiome.^21^

QS autoinducers can be species-specific, meaning that a single bacterial species produces and detects a particular autoinducer. Other autoinducers are more “universal”; they are produced and detected by many bacterial species. Autoinducer-2 (AI-2) is among the latter class and is produced by the LuxS synthase, which is widely conserved in bacteria.^22–24^ AI-2 consists of a set of interconverting compounds derived from 4,5-dihydroxy-2,3-pentanedione (DPD, **1** in **Figure 1A**), an intermediate in the *S*-adenosylmethionine (SAM)-dependent methylation recycling pathway.^22^ Following release of an activated methyl group from SAM to an acceptor substrate, the byproduct, *S*-adenosylhomocysteine (SAH), undergoes a two-step transformation to DPD: SAH is converted to *S-*ribosylhomocysteine (SRH) by the enzyme Pfs, followed by further processing by LuxS, yielding homocysteine and compound **1**. In aqueous solution, compound **1** rapidly interconverts between linear, cyclic, and hydrated forms that coexist in equilibrium.^25–28^ The active AI-2 moiety that is recognized depends on the particular bacterial receptor, as well as the chemical environment. The boron-rich marine environment promotes formation of a boron adduct (2*S*,4*S*)-2-methyl-2,3,3,4-tetrahydroxytetrahydrofuran-borate (*S*-THMF-borate, **2**), which is the active AI-2 recognized by the LuxP receptor in *Vibrio* spp.^29^ In the absence of boron, compound **1** rearranges to (2*R*,4*S*)-2-methyl-2,3,3,4-tetrahydroxytetrahydrofuran (*R*-THMF, **3**), which is the active AI-2 recognized by the LsrB receptor in enteric bacteria.^27^ LsrB, first identified in *Salmonella enterica ssp.* typhimurium, is structurally similar to LuxP and occurs in *Escherichia coli* as well as some members of the *Clostridiaceae* and *Bacillacaeae* families.^30, 31^ The ubiquitous pathogen, *Pseudomonas aeruginosa*, encodes a receptor with a dCache_1 domain, a substructure that binds AI-2 in LsrB and LuxP.^32^ The *P. aeruginosa* receptor is reported to bind AI-2. Over 1,500 transmembrane proteins harboring the dCache_1 domain are known among bacteria and archaea, providing opportunities for discovery of potentially new AI-2 structures and functions.^32, 33^

**Figure 1.**
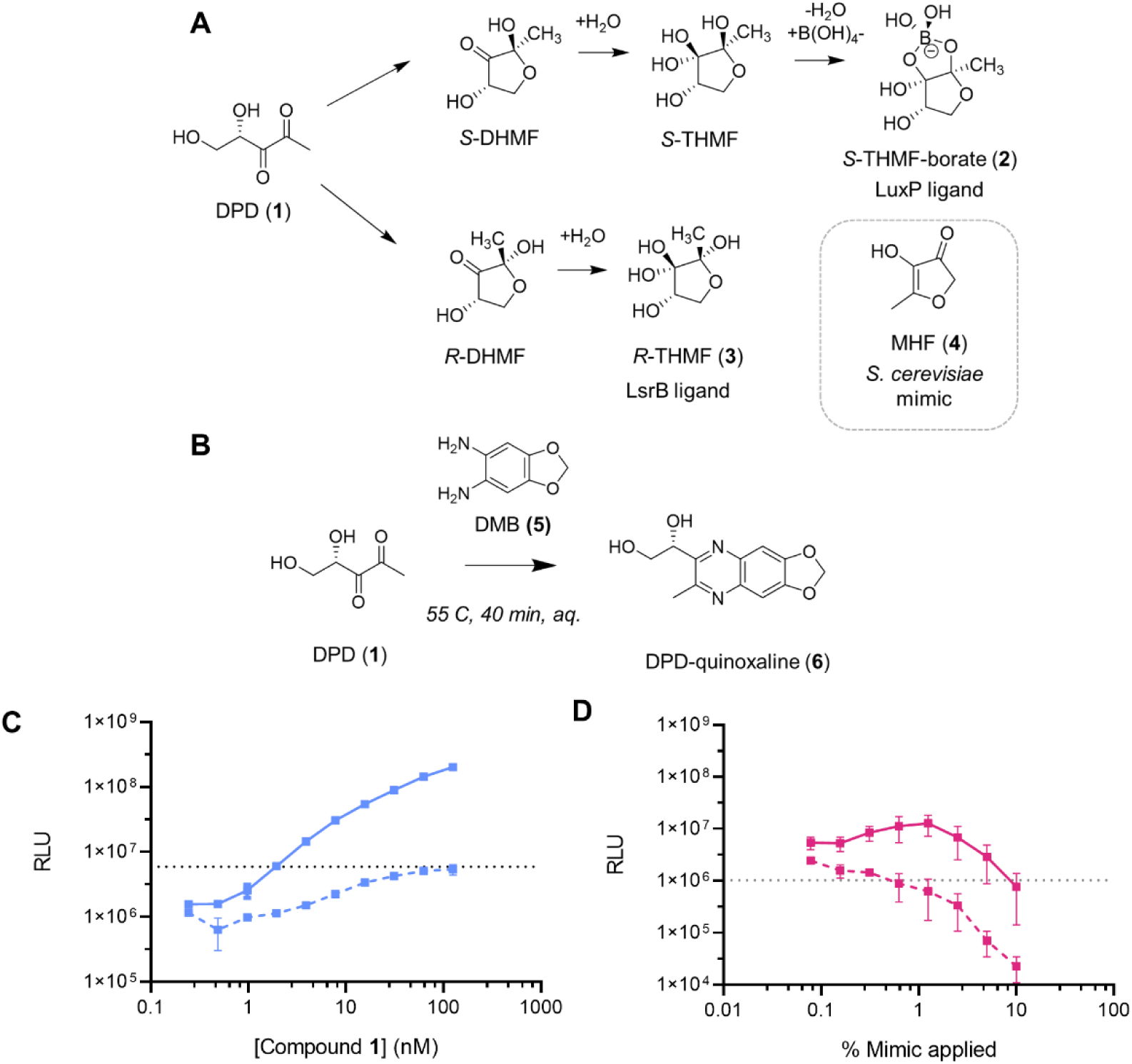
Derivatization of α-diketone compounds as a chemical screening strategy to identify the mammalian AI-2 mimic. A) Diagram showing the structure of the AI-2 precursor DPD (**1)**, known interconverting AI-2 moieties (**2**) and (**3**), and the *S. cerevisiae* AI-2 mimic MHF (**4**). B) The *o-*diaminobenzene derivatization reagent (**5**) and the reaction scheme used for derivatizing compound **1** to form the functionalized quinoxaline product (**6)**. C) Monitoring reaction completion by measuring the activity of compound **1** via light output from the *V. harveyi* TL-26 bioassay. Mixtures containing a 5 µM solution of compound **1** incubated with either control solution (solid line) or 3 mM compound **5** (dashed line) were applied in serial dilution to *V. harveyi* TL-26. The concentrations designated on the x-axis are relative to the initial concentration of compound **1** in the control reaction. D) Light output from *V. harveyi* TL-26 in response to 50 µl of culture fluids from Caco-2 cells treated 1:1 (v/v) with either control solution (solid line) or 3 mM compound **5** (dashed line). At high concentrations, both the control and reacted material cause declines in light output from the reporter, presumably due to the presence of inhibitors/toxic compounds. In C and D, RLU denotes relative light units, which are bioluminescence/OD_600_, the dotted lines show the baseline level bioluminescence in the assay, and error bars represent standard deviations of biological replicates, *n*=3.

Many strains of gut-associated bacteria across the Bacteroidetes, Firmicutes, and Proteobacteria phyla possess the AI-2 biosynthetic enzyme LuxS.^34–36^ The yeast *Saccharomyces cerevisiae* produces an AI-2 mimic, 4-hydroxy-5-methylfuran-3(2H)-one (MHF, compound **4**), that agonizes LuxP.^37^ The yeast MHF-synthase Cff1p has homologs across archaeal, bacterial, and fungal genomes. The human host also participates in AI-2-driven communication. Specifically, in response to the secreted bacterial cytolytic toxin aerolysin, tight-junction disruption, or nutritional stress, mammalian intestinal epithelial cells produce an AI-2 mimic, i.e., a compound harboring AI-2 activity.^38^ Prior to this report, the identity of the mammalian AI-2 mimic had not been reported, but the active compound appeared not to be one of the previously identified AI-2 structures for the following reasons: First, eukaryotic genomes do not possess *luxS*. Second, supplementation of SRH to mammalian cell culture did not promote increased AI-2 mimic production, indicating that epithelial cells cannot convert SRH to compound **1**. Third, HPLC fractionation of epithelial cell culture fluids showed that the mammalian AI-2 mimic activity did not elute with compound **1**, indicating that the compounds are not identical.

In this study, we developed a reactivity-based metabolomics workflow based on chemical derivatization that allowed us to capture a chemically unstable mammalian AI-2 mimic, thus enabling its identification and characterization. We identify the metabolite L-xylosone as an AI-2 mimic from human Caco-2 epithelial cells. We find that L-xylosone is recognized by the bacterial LuxP AI-2 receptor with micromolar affinity, whereas the D-enantiomer is not. Examination of compounds with structural features similar to L-xylosone revealed that L-xylulose, a primary metabolite in the glucuronate-xylulose (GX) pathway, also harbors AI-2 activity. Our identification of two mammalian-produced autoinducer mimics begins to define the chemical lexicon employed in cross-domain host-bacterial communication.

## Results

### Development of an α-diketone-based derivatization strategy to trap a mammalian AI-2 mimic

AI-2 has posed significant challenges for spectroscopic detection and characterization. As noted above, the AI-2 precursor (**1**) exists in an equilibrium mixture of linear and cyclic forms that undergo subsequent hydration events.^25–28^ There is no UV chromophore in the set of molecules comprising AI-2 and they ionize poorly using standard MS techniques. For these reasons, traditional activity-guided compound isolation did not reveal the structures of AI-2; rather, crystallization and structure elucidation of the LuxP-ligand and LsrB-ligand complexes provided the identities of the active AI-2s.^27, 29^ Likewise, the mammalian AI-2 mimic did not yield to traditional purification and characterization methods, and its structure has remained unknown.

Previous analytical strategies to detect and quantify compound **1** in biological samples have hinged on derivatization, primarily relying on an *o-*diaminobenzene tag that reacts specifically with the α-diketone moiety of compound **1**.^39–42^ The resulting quinoxaline product provides sensitive and reliable UV detection, as well as a specific *m/z* signature. We reasoned that the mammalian AI-2 mimic must be structurally similar to compound **1** and would also contain an α-diketone functional group. Leveraging this reactivity for tagging and detection could make the mammalian AI-2 mimic amenable to LC-MS-based metabolomics.

We evaluated multiple *o-*diaminobenzene containing derivatization agents and selected 1,2-diamino-4,5-methylenedioxybenzene (DMB, **5**, **Figure 1B**) for our initial analyses as the derivatization reaction proceeds in complex aqueous biological matrices and because **5** is commercially available.^43^ The reaction between compounds **1** and **5** was optimized, and quinoxaline product formation was monitored by LC-MS (compound **6**, **Figure S1**). Reaction completion was assessed by applying the reaction mixture to a *Vibrio harveyi* strain that reports on AI-2 activity. This *V. harveyi* strain, called TL26, carries a *luxS* deletion (Δ*luxS*) and is therefore incapable of AI-2 production.^44^ *V. harveyi* TL26 emits bioluminescence only when supplied with exogenous AI-2. Reacting compound **1** with excess compound **5** eliminated all AI-2 activity in the sample as judged by the *V. harveyi* TL26 bioassay (**Figure 1C**), suggesting that AI-2 had been completely derivatized under these conditions. The presence of compound **5** is not toxic, nor does it interfere with bioluminescence. Specifically, compared to **1** alone, addition of compounds **1** and **5** together at the start of the bioassay did not decrease *V. harveyi* TL26 light output (**Figure S1**).

Preparation of the mammalian AI-2 mimic for analyses followed a previously reported procedure. Briefly, medium from cultured Caco-2 cells was replaced with phosphate buffered saline (PBS), the cells were incubated for 48 h, and cell-free culture fluids harvested. Treatment of Caco-2 cells with PBS drives AI-2 mimic production while not affecting Caco-2 cell viability.^38^ We reacted multiple such preparations of the mammalian AI-2 mimic with **5**. When we applied the derivatization reaction mixtures to *V. harveyi* TL26 all bioassay activity was eliminated (**Figure 1D**). To confirm the selectivity of this reactivity-based chemical screen, we incubated compound **5** with MHF (compound **4**), the yeast AI-2 mimic which, notably, lacks an α-diketo-moiety. No fluorescence emission indicative of a quinoxaline derivative was observed in reactions between compounds **4** and **5 (Figure S2**).^43^ When this reaction mixture was applied to *V. harveyi* TL26, light output was nearly identical to the untreated sample. Indeed, even a reaction with 100-fold molar excess of **5** did not eliminate activity of compound **4** in the bioassay (**Figure S2**). The reactivity profile of **5**, coupled with its ability to inactivate mammalian AI-2 mimic activity, support our hypothesis that an α-diketone-containing compound primarily accounts for the mammalian AI-2 mimic activity.

### Identification of xylosone as a mammalian AI-2 mimic

We employed a reactivity-guided metabolomics screen to identify the mammalian AI-2 mimic **(Figure 2A)**. Samples for our chemical screen were prepared as follows: Preparations of mammalian AI-2 mimic were incubated with compound **5** at 55°C for 40 min and analyzed by LC-MS. As a control, compound **5** was likewise incubated in PBS followed by LC-MS analysis. The initial dataset revealed 2,897 differentially abundant features (**Figure S3**). To identify biologically relevant signals, we applied stringent filters to limit our focus to metabolites with an LC-MS intensity threshold of 10,000, exhibiting at least a 10-fold increase compared to the control, and possessing *q* values ≤ 0.01 and *m/z >*175, all features of compounds that could contain the quinoxaline core resulting from reaction with **5**.

**Figure 2.**
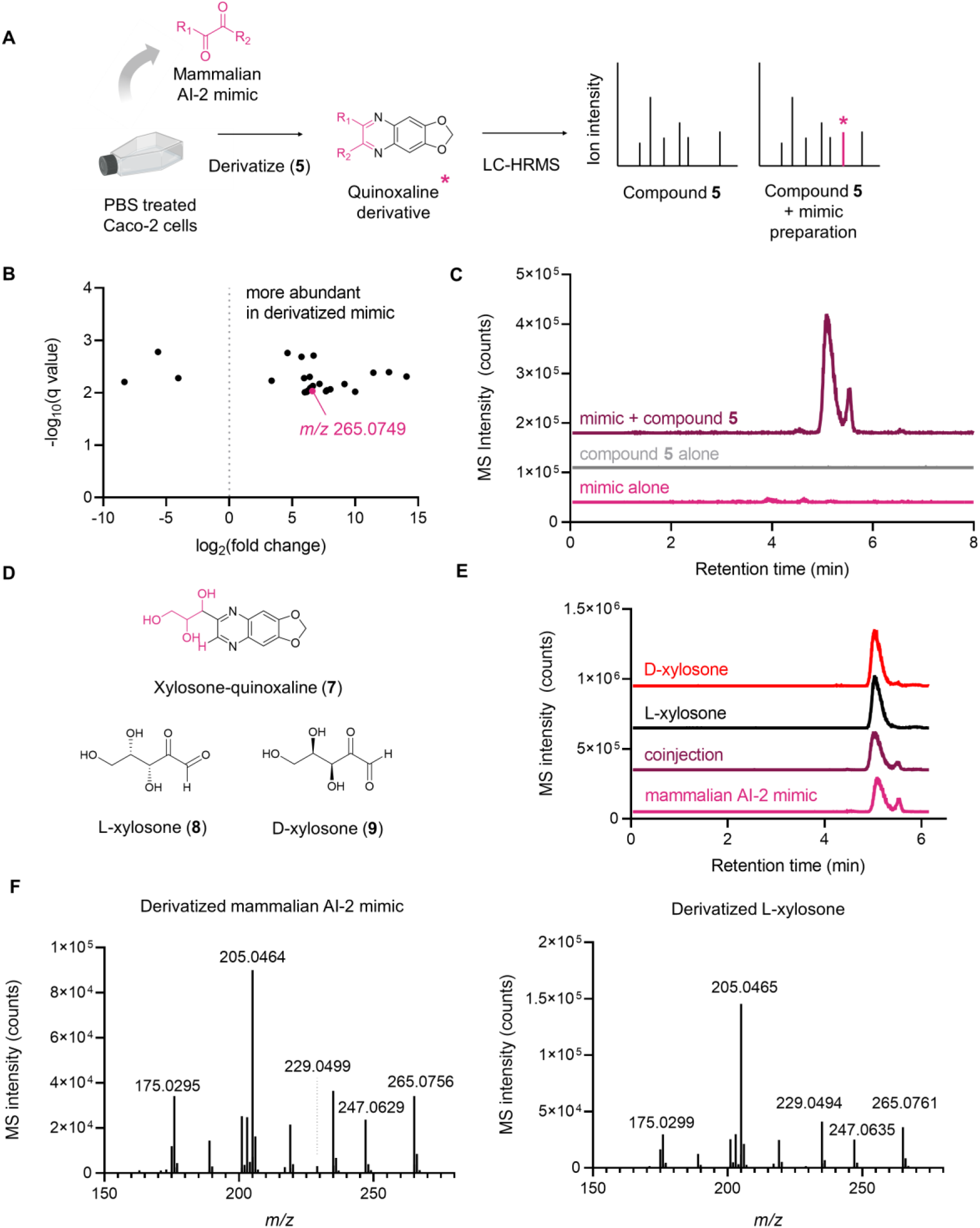
Comparative LC-MS analysis of mammalian AI-2 mimic samples derivatized with **5** identifies xylosone (**8** and **9**). A) Reactivity-based screening workflow. B) Volcano plot in which each data point represents a distinct molecular species detected by LC-MS. Features were filtered to include only those with *m/z* >175, a minimum MS intensity threshold of 10,000, >10-fold changes compared to the control, and *p* values < 0.001. The observed ion for xylosone-quinoxaline (**7**, see panel D) is highlighted in pink. C) Extracted ion chromatogram (EIC) spectra for *m/z* 265.0819 from mammalian AI-2 mimic samples treated with (maroon) or without (pink) **5**, and of compound **5** alone (gray). D) Proposed chemical structure of the xylosone-quinoxaline product, compound **7** and L- and D-xylosone (compounds **8** and **9**, respectively). E) EIC spectra (*m/z* 265.0819) of mammalian AI-2 mimic (magenta), synthetic **8** (black), or synthetic **9** (red), each derivatized with **5**. Also shown is a 1:1 v/v coinjection of **5**-derivatized mammalian AI-2 mimic and compound **8**. F) MS/MS spectra (*m/z* 265.0819) of **5**-derivatized AI-2 mimic and compound **8**. Collision energy = 20 eV.

We mined the resulting 25 molecular features (**Table S1**) for predicted *m/z* corresponding to quinoxaline derivatives of previously reported α-diketone metabolites.^45^ This strategy revealed a unique mass feature corresponding to *m/z* 265.0749 (**Figure 2B**), detected as two resolved peaks, one major (retention time = 5.1 min) and one minor (retention time = 5.5 min). This species was present in mammalian AI-2 mimic preparations derivatized by compound **5**, but not in samples that had not been subjected to derivatization (**Figure 2C**). The molecular formula of the putative α-diketone-containing metabolite was determined to be C_5_H_8_O_5_, matching that of xylosone, a rare sugar that is new to human cells and can exist as L- or D-enantiomers (**8** and **9**, **Figure 2D**).

To verify xylosone as the active component and differentiate between the two enantiomers, we developed a synthetic route to obtain pure **8** and **9** (**Figure 3**). To obtain **8**, we began with L-xylose (**10**), converted it to the acetyl-protected, cyclic form (**11**) and installed the (*1R*)-thioketal regio- and diastereoselectively by using boron trifluoride etherate and thiophenol (**12**).^46^ The remaining acetate groups in **12** were then deprotected, followed by acetonide protection of vicinal diols in **13** that resulted in the formation of two regio-isomers **14b** and **14a** in an 8:1 ratio and an 86% combined yield.^47^ The desired product **14a** is the minor regio-isomer formed during the acetonide protection, which was separated from **14b** and subjected to Swern oxidation using oxalyl chloride, DMSO, and triethylamine to furnish ketone **15** in 75% yield. Global deprotection of the thioketal and acetonide groups in **15** using *N*-Bromosuccinimide afforded L-xylosone (**8**) in 64% yield. This route also allows for the preparation of D-xylosone (**9**) from D-xylose.

**Figure 3.**
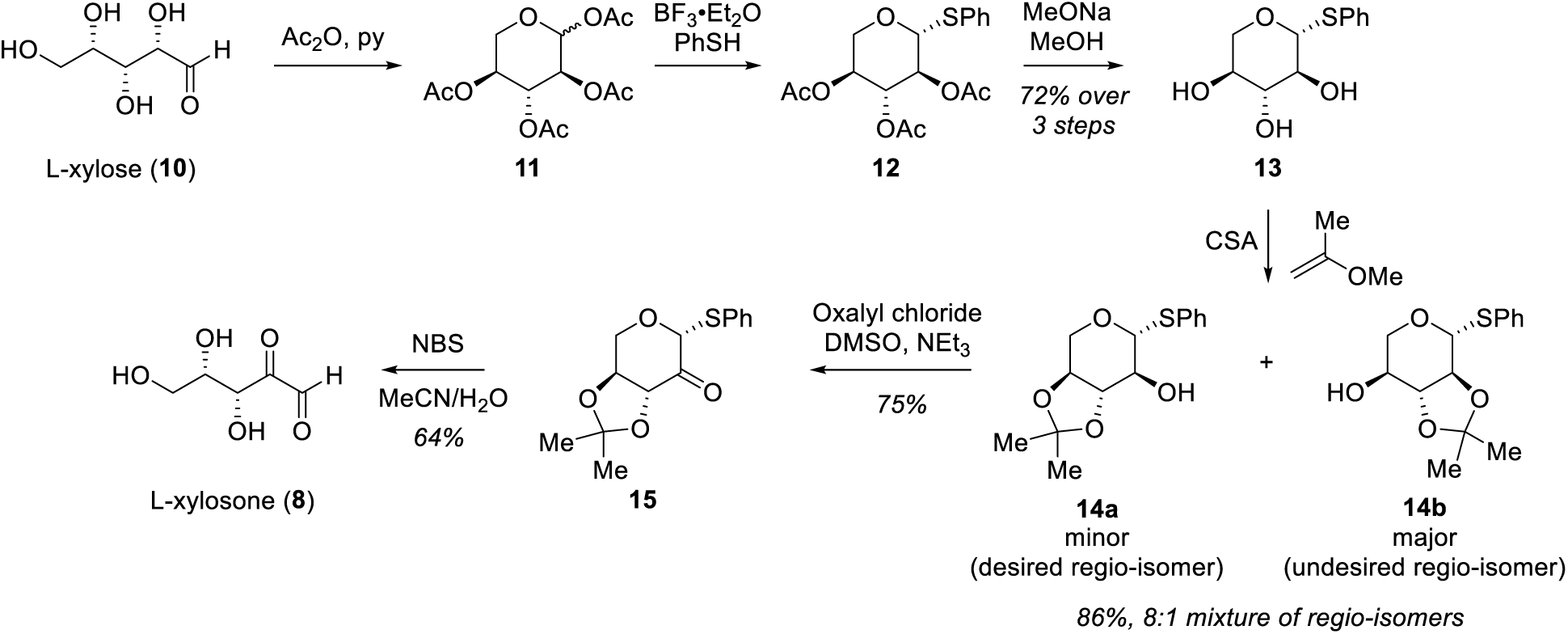
Synthetic scheme used to access enantiopure L- and D-xylosone. The scheme is shown for L-xylosone (**8**) starting with L-xylose, but the identical approach was used to obtain D-xylosone (**9**) starting from D-xylose. L-xylose (**10**) is first converted into **11** using a combination of acetic anhydride (Ac_2_O) and pyridine (py). The thioketal in **12** was then installed regio- and diastereoselectively by activating the anomeric acetate group in **11** using boron trifluoride etherate (BF_3_•Et_2_O) and thiophenol (PhSH). Deprotection with sodium methoxide (MeONa) in methanol yielded **13**. The vicinal diols in **13** were then acetonide-protected using camphor sulfonic acid (CSA), resulting in the formation of two regio-isomers **14b** and **14a** in 8:1 ratio and 86% combined yield. The desired product **14a** is the minor regio-isomer formed during the acetonide protection, which was separated from **14b** and subjected to Swern oxidation conditions using oxalyl chloride, DMSO and triethylamine (NEt_3_) to furnish ketone **15** in 75% yield. Global deprotection of the thioketal and acetonide groups in using *N*-Bromosuccinimide (NBS) in acetonitrile/water afforded L-xylosone (**8**) in 64% yield.

With the two enantiomers in hand, we prepared the quinoxaline derivatives of synthetic L-xylosone and D-xylosone, which exhibited identical chromatographic properties and were not separable by HPLC with a retention time of 5.1 min. Importantly, this elution profile matched the retention time of the derivatized mammalian AI-2 mimic (**Figure 2E**). Coinjection of the derivatized synthetic material and the mammalian AI-2 mimic also showed identical retention times (**Figure 2E**). Finally, MS/MS analysis confirmed that the derivatized synthetic material was indistinguishable from mammalian-derived xylosone-quinoxaline using a range of collision energies (20 to 50 eV) (**Figure 2F**, **Figure S4, Tables S2, S3**). These data verify xylosone as an active AI-2 mimic.

### Verification that L-xylosone possesses AI-2 activity

We next assessed each xylosone enantiomer for AI-2 mimic activity using the *V. harveyi* TL26 bioassay (see **Figures S5-S9** for quantitation methods).^48^ L-xylosone (**8**) was active with a half-maximal effective concentration (EC_50_) of ∼2 µM, while D-xylosone (**9**) was inactive (**Figure 4A, Figure S16**). The stereochemical selectivity for the L-xylosone enantiomer aligns with prior findings from the crystal structure of the LuxP-AI-2 complex.^29^ The hydroxyl group on the C-4 of **2** forms two hydrogen bonds with LuxP residues Trp82 and Gln77. Inversion from the *S-* to the *R-*configuration renders the hydroxyl group inaccessible for hydrogen bonding.^49^ Compound **8** retains the C-4 configuration that is required for binding to the LuxP receptor. The human intestinal pathogen and quorum-sensing bacterium *Vibrio cholerae* detects AI-2 to control traits including virulence and biofilm formaiton.^50–53^ A *V. cholerae* AI-2 reporter strain^53^ responded to exogenous compound **8** with an EC_50_ of ∼8.3 µM, comparable to the measured EC_50_ of ∼ 1 µM for the native ligand, compound **1** (**Figure S10**).

**Figure 4.**
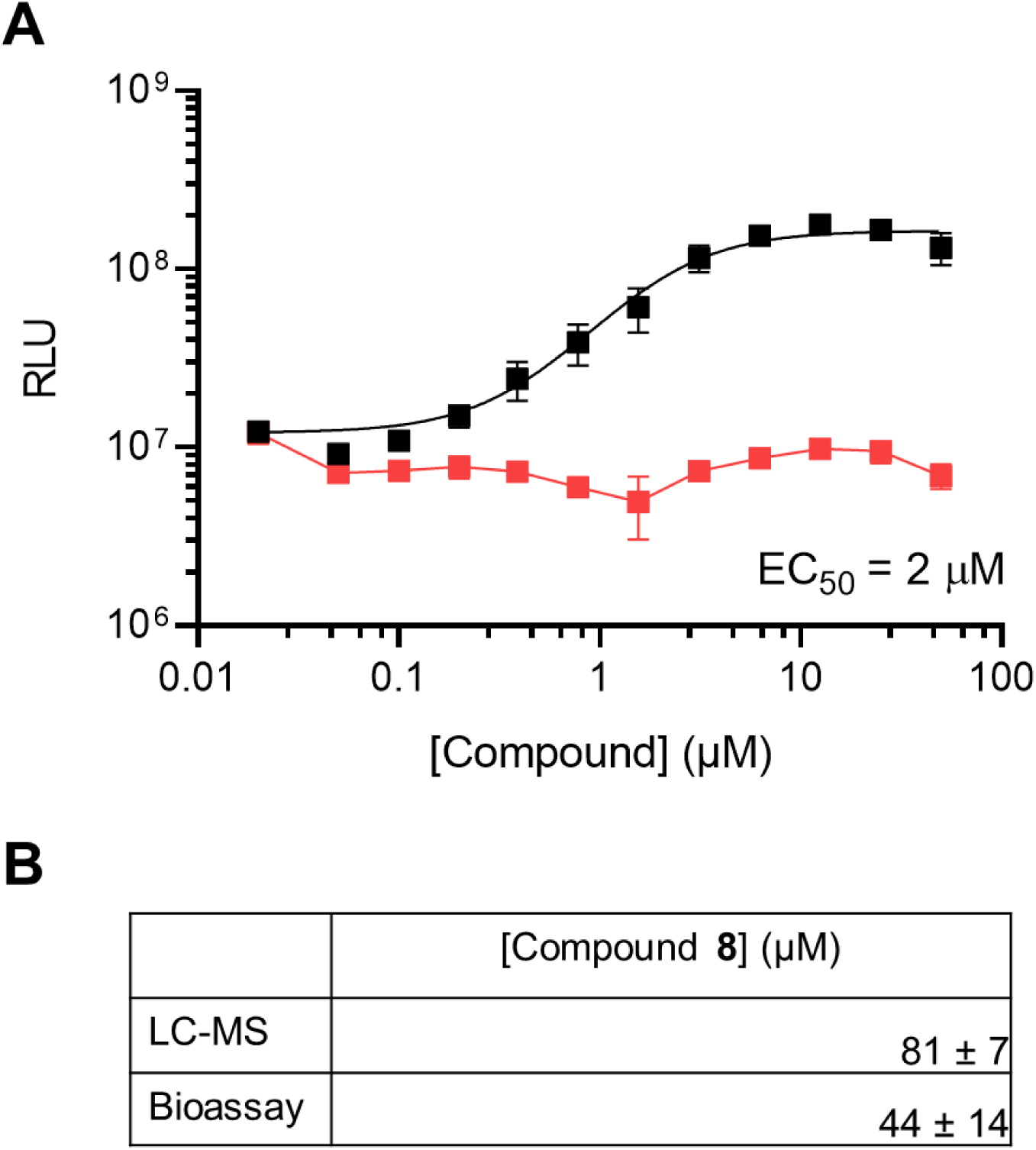
L-xylosone (**8**) has AI-2 activity, while D-xylosone (**9**) does not. A) Light output from the *V. harveyi* TL-26 bioassay in response to indicated amounts of **8** (black) and **9** (red). The EC_50_ for **8** is shown. Error bars represent standard deviations of technical replicates, *n* = 3. B) Quantitation of **8** from Caco-2 AI-2 mimic preparations from 2,000,000 cells grown for 2 days. Concentrations are calculated from LC-MS measurements and the *V. harveyi* TL-26 bioassay. Data are reported as averages ± standard deviations of biological replicates, *n*=3.

Analogous to other AI-2s, compound **8** likely cyclizes to form a furanose that mimics compound **2** in the binding pocket of LuxP. Indeed, the ^13^C NMR spectrum of compound **8** displays 6 major signals suggestive of the formation of at least 3 acetal and/or hemiacetal moieties of the α-ketoaldehyde in xylosone (**Figure S11**). We conducted boron binding studies with **8** with the rationale that cyclization and subsequent hydration of the C-3 carbonyl of **2** could lead to formation of a borate diester that drives maximal AI-2 activity in the *V. harveyi* bioassay.^27–29^ Similarly, addition of boric acid to the medium is required to achieve high potency of compound **8** in the bioassay (**Figure S12A, B**), supporting the notion that L-xylosone complexes with borate across a cis-diol. Borate esters of 1,3-diols and 1,2-diols display characteristic chemical shifts, allowing distinction between these moieties.^54^ Surprisingly, however, no ^11^B or ^13^C NMR signals indicative of borate-diester formation were detected when borate was incubated with compound **8** (**Figures S13, S14**). It is possible that formation of the borate diester complex of **8** only occurs in the LuxP binding pocket.

We used two methods to quantify the amount of xylosone present in mammalian culture fluids. First, using the *V. harveyi* TL26 bioassay, we estimated concentrations based on measurements of activity from known quantities of pure **8**. Second, mammalian AI-2 mimic samples were derivatized with **5**; then, using LC-MS we determined the concentration from a standard curve generated from known amounts of **8** derivatized with **5**. The two methods yielded similar inferred concentrations: After 2 days of growth in PBS, Caco-2 cells produced 44 ± 14 µM and 81 ± 7 µM AI-2 mimic as determined by bioassay and LC-MS, respectively (**Figure 4B**). For comparison, recent GC-MS based quantitation of compound **1** levels in the cecal contents of specific pathogen free mice ranged from 0.07 – 0.21 µM^41^, consistent with reported K_d_ values for LsrB receptors from various bacterial species.^31^ Together, these data confirm L-xylosone as the active AI-2 mimic and provide production titers that appear physiologically relevant and significantly higher than the EC_50_ values determined in the *V. harveyi* TL26 and *V. cholerae* bioassays (**Figure 4A**).

### L-xylulose, a compound structurally related to L-xylosone possesses AI-2 activity

We tested two commercially available compounds, xylose and xylulose, with structural features similar to compound **8** for AI-2 mimic activity. These compounds differ from **8** only in the oxidation state of a single carbon atom. Neither the naturally occurring D-xylose (**16**) nor L-xylose (**10**) possessed detectable AI-2 activity (**Figure S15**). Likewise, D-xylulose (**17**) is also inactive, consistent with prior reports (**Figure 5A**).^49^ By contrast, L-xylulose (**18**) was active, with an EC_50_ ∼6.0 µM (**Figure 5A, Figure S17**). L- and D-xylulose are naturally occurring sugars generated as intermediates in the glucuronate-xylulose pathway leading to D-xylulose-5-phosphate. Analogous to other LuxP ligands, addition of boric acid to the medium is required for optimal activity (**Figure S12C**) and a small amount of complex could be detected by ^11^B NMR (**Figure S16;** chemical shift δ 10.6 ppm), consistent with previously reported values.^55^ With respect to detection by the *V. harveyi* quorum-sensing apparatus, compound **1** is most potent, followed by compounds **8** and then **18** (**Figure S17**).

**Figure 5.**
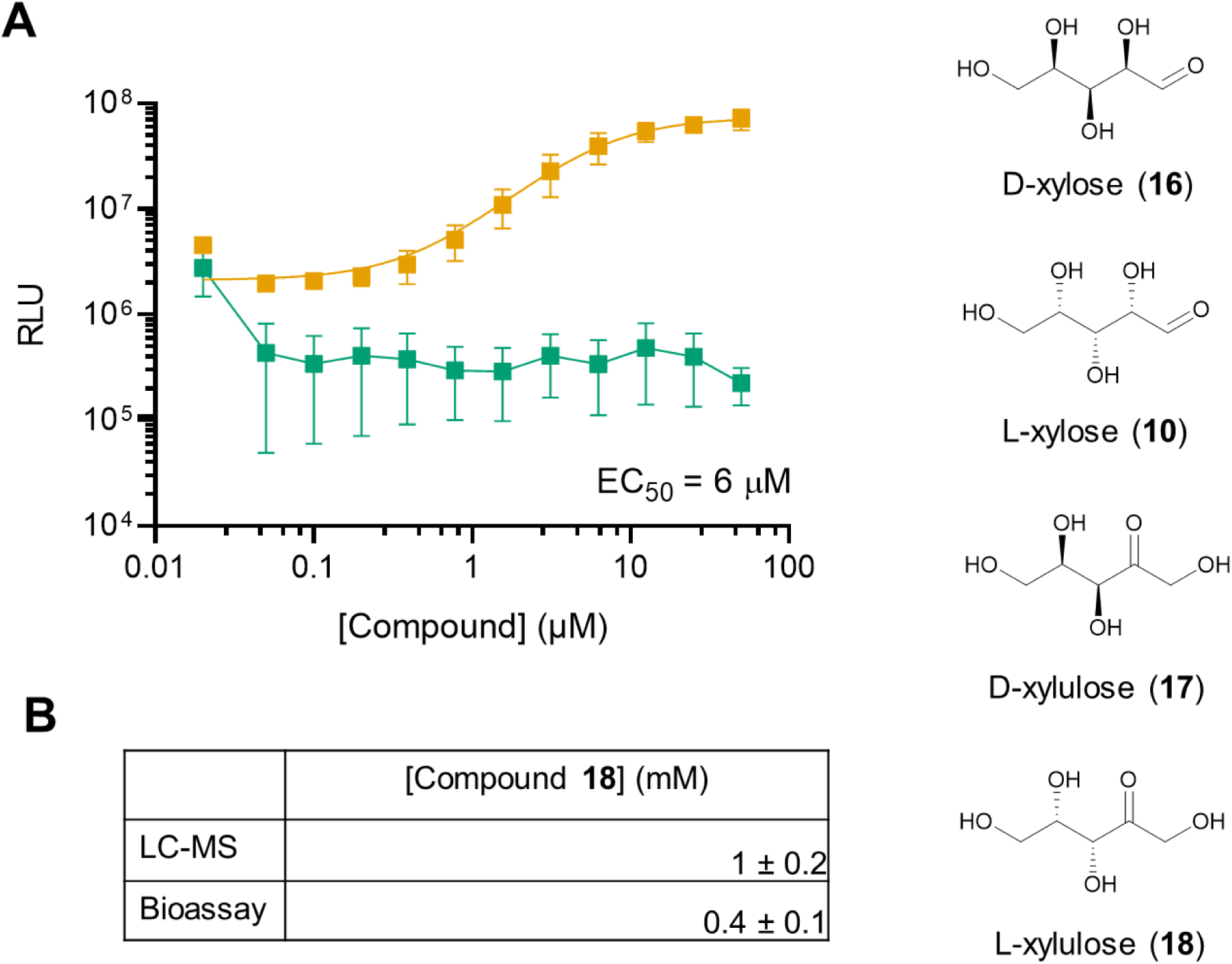
The structurally related ketopentose sugar L-xylulose (**18**) possesses AI-2 activity. A) Light output from the *V. harveyi* TL-26 bioassay in response to indicated amounts of **17** (green) and **18** (orange). The EC_50_ for **18** is provided. Error bars represent standard deviations of technical replicates, *n* = 3. B) Quantitation of **18** levels in Caco-2 AI-2 mimic preparations from 2,000,000 cells grown for 2 days. Concentrations are calculated from LC-MS measurements and the *V. harveyi* TL-26 bioassay. Data are reported as averages ± standard deviations of biological replicates, *n* = 3.

To determine the production titers of **18** by Caco-2 cells under our experimental conditions, we derivatized the cell-free fluids with *N*,*N*-diphenylhydrazine and used LC-MS to quantify the hydrazone derivative (**Figure 5B**).^48^ We also estimated the concentration of compound **18** present in mammalian mimic preparations by comparison to the *V. harveyi* TL26 bioassay activity generated from known amounts of compound **18.** These assays yielded a concentration of 0.4 ± 0.1 mM (bioassay) and 1 ± 0.2 mM (LC-MS). We suspect that these values are overestimates because the bioassay output will be the sum of activity from **8** and **18** and our LCMS method does not distinguish between the hydrazone derivative of xylulose and the isobaric hydrazone derivative of xylose.

## Discussion

The microbiome-mucosal interface is a crucial site for regulation of intestinal function. On the bacterial side, microbial products such as short-chain fatty acids,^56^ secondary bile acids,^57, 58^ and tryptophan metabolites^59, 60^ function to maintain gut epithelial integrity. On the host side, intestinal epithelial cells detect pathogen-associated molecular patterns (PAMPs), including flagellin and lipopolysaccharide, on the outer surfaces of bacterial cells and they relay the garnered information to intestinal immune cells to launch defense mechanisms and drive immune tolerance.^61^ Mammalian metabolites, including antimicrobial peptides, cytokines, and secreted IgA antibodies shape microbiome composition.^62^ Adding to these bidirectional chemical interactions, we previously reported a mammalian AI-2 mimic of unknown structure that is detected by bacterial cells.^38^ Here, our studies reveal that the new human metabolite L-xylosone functions as a QS autoinducer to which bacteria respond. Prior to this report, the only known role of xylosone was its involvement in non-enzymatic modifications of proteins to form advanced glycation end products (AGEs).^63–65^ Our work demonstrates that L-xylosone functions as a QS signal that bacteria detect; we do not yet know whether L-xylosone has additional metabolic and/or signaling roles affecting the mammalian host.

The most highly studied in vitro route to xylosone formation is from the spontaneous, oxidative degradation of ascorbic acid to dehydroascorbic acid.^66, 67^ Under neutral or alkaline conditions, dehydroascorbic acid undergoes hydrolytic cleavage of the lactone ring to yield 2,3 diketo-L-gluconic acid, which subsequently decarboxylates to L-xylosone, which is highly reactive. Humans are incapable of synthesizing ascorbic acid and therefore obtain it from diet. Although often a common component of mammalian cell culture medium, we note that ascorbic acid is not present in media used here. Indeed, we were unable to detect ascorbic acid in our media nor in mammalian AI-2 mimic preparations using LC-MS. Thus, L-xylosone must originate from a non-ascorbic acid route. Some eukaryotes, such as wood-degrading fungi, harbor pyranose 2-oxidase (P2Ox) which can produce xylosone.^68^ P2Ox enzymes are flavoenzymes that, in the presence of oxygen, catalyze the regioselective C-2 oxidation of D-glucose, D-xylose, and various other mono- and di-saccharides to yield the corresponding dicarbonyl sugars and hydrogen peroxide.^69–71^ P2Ox enzymes belong to the glucose-methanol-choline (GMC) oxidoreductase superfamily.^70^ A search of the InterPro database reveals two mammalian genes encoding proteins belonging to this class of enzymes: CHDH, encoding a choline dehydrogenase, and B4DMQ4, encoding a gene of unknown function similar to CHDH. Conceivably, one or both of these enzymes could catalyze C-1 oxidation of L-xylulose to produce L-xylosone. Lastly, it is also possible that L-xylosone production occurs spontaneously as is the case for other dicarbonyl compounds that are formed through dehydration and enolization of monosaccharides.

We also demonstrated that L-xylulose has AI-2 mimic activity. More is known about L-xylulose than L-xylosone in human metabolism. L-xylulose is a minor sugar produced in the glucuronate degradation pathway, an alternate pathway for glucose-6-phosphate oxidation.^72^ Specifically, L-xylulose is produced by decarboxylation of β-keto-L-gulonate by the C11orf54 protein.^73^ Next, L-xylulose is reduced to xylitol in a reversable reaction catalyzed by the dicarbonyl/l-xylulose reductase DCXR. Mutations in DCXR result in excretion of L-xylulose, the hallmark of the metabolic disorder called pentosuria.^74^ DCXR also functions as a diacetyl reductase that detoxifies highly reactive α-dicarbonyl compounds.^75, 76^ Indeed, DCXR deficient mice are more vulnerable to protein damage following diacetyl-induced cytotoxicity than are mice that are wildtype for DCXR.^77^ One possibility is that alterations in DCXR activity, triggered by starvation or tight junction disruption, decrease DCXR diacetyl reductase activity and drive the accumulation and secretion of L-xylulose and L-xylosone.

We cannot exclude the possibility of additional mammalian AI-2 mimics, as our study specifically targeted α-diketone containing compounds. Indeed, L-xylulose is not derivatized in our chemical trapping strategy so it was not identified in our dataset, yet we show it has AI-2 mimic activity. To our knowledge, there is no Caco-2 cell line incapable of producing L-xylulose and L-xylosone, which limits our ability to probe whether multiple AI-2 mimics exist. Other AI-2 mimics could also be produced when human cells are cultured under conditions that differ from those employed here. In this work, we induced epithelial cells to produce L-xylosone using nutrient deprivation stress. Earlier, we showed that AI-2 mimic production occurs following epithelial cell exposure to toxins or tight junction disruption. In those cases, we have not confirmed that L-xylosone or L-xylulose is the compound produced that harbors the AI-2 mimic activity.

Our findings highlight the utility of reactivity-guided metabolomics approaches to capture chemically unstable metabolites of interest. Reactivity-based screening approaches have been previously used to target labile functional groups such as ⍺,β-unsaturated carbonyls,^78, 79^ isonitriles,^80^ epoxides,^81^ and diazo^82^ groups. Using our approach to discover α-diketone containing compounds, we detected the rare metabolite L-xylosone, which was not previously identified in the human metabolome database. Mammalian α-diketone containing compounds of intense research focus have historically centered on methylglyoxal, glyoxal, and 3-deoxyglucosone for their roles in the formation of AGEs in aging tissues.^45, 83–85^ We speculate that there other highly reactive α-diketone containing compounds exist that are likewise absent from existing metabolomic datasets that possess novel and potentially fascinating biological roles.

## Experimental Methods

### Mammalian AI-2 mimic production and preparation for analysis

Caco-2 cells were grown at 37°C and 5% CO_2_. For maintenance, Caco-2 cells were grown in 1X DMEM (Gibco), 20% FBS (Corning), 1X Penstrep (Sigma), and 1x Plasmocin prophylactic (Invivogen). For mammalian AI-2 mimic production, Caco-2 cells were grown to confluence, washed twice with 1X Dulbecco’s PBS (Gibco), and detached from tissue culture plates using trypsin-EDTA (Corning). These cells were collected, washed once with maintenance medium to inactivate the trypsin, washed twice with 1X PBS, and subsequently placed in 5 mL of 1X PBS at a cell density of 2,000,000 cells/mL for 48 h. Cell-free culture fluids were harvested after centrifugation at 1,500 rpm for 5 min followed by filtration through a 0.22 µm filter. We call these samples mammalian AI-2 mimic preparations throughout this work. These preparations were stored at 4°C prior to use.

### AI-2 bioassay

*V. harveyi* TL-26 was grown overnight in LM medium at 30°C and diluted 1:1000 into AB medium supplemented with 0.1 mM boric acid. *V. cholerae* AB_Vc_542 (Δ*cqsS* Δ*luxQ* Δ*vpsS* Δ*cqsR* Δ*vc1807*::*PluxC-luxCDABE*::Spec^R^) was grown overnight in LB medium at 30°C and diluted 1:5000 into LB medium supplemented with 0.1 mM boric acid. The diluted cultures were aliquoted into wells of black-sided, clear-bottom 96-well plates (Corning). All compounds, derivatization reaction mixtures, and mammalian AI-2 mimic preparations were added at the indicated amounts. Plates were incubated at 30°C with shaking for 6 h, followed by bioluminescence measurements using an Envision plate reader (PerkinElmer).

### General LC-MS Methods

Low resolution data for optimization of compound **5** derivatizations were acquired using an Agilent MSD iQ System coupled to an Agilent 1290 Infinity II HPLC. Samples were separated on an Agilent Eclipse C18 1.8 µm (2.1 x 50 mm) column with an injection size of 2 µL. The mobile phase was a water-acetonitrile (MeCN) gradient containing 0.1% formic acid. Chromatography was performed as follows: 0-2 min 5% MeCN, 2-10 min 5-95% MeCN, 10-12 min 95% MeCN. The LC-MS iQ was carried out in positive mode scanning between m/z 100-1000. Source parameters for LC-MS acquisition were as follows: gas temperature 325°C, gas flow 11 (L/min), capillary voltage 3500 V. Data were processed using Agilent OpenLAB CDS. Peaks were extracted by *m/z,* quantified by area under the curve. High resolution LC-MS data for the metabolomics workflow and quantitation of compounds **8** and **18** in mammalian AI-2 mimic samples were acquired on an Agilent 6546 LC-QTOF 1290 LC system. Samples were separated on a Phenomenex Polar RP 2.5 µm (100 x 3 mm) column with an injection size of 5 µL. The mobile phase was a water-MeCN gradient containing 0.1% formic acid. Chromatography was performed as follows: 0-2 min 5% MeCN, 2-10 min 5-95% MeCN, 10-11 min 95% MeCN, equilibrate to 5% MeCN. MS1 acquisition was carried out in positive mode scanning from 100-500 *m/z*. MS2 acquisition was carried out in positive mode scanning from 100-600 *m/z* with fixed collision energies at 20, 25, 55 eV. The MS1 scan rate was 3 spectra/sec and the MS/MS scan rate was 1 spectra/sec. Source parameters for both MS1 and MS2 acquisition were as follows: gas temperature 275°C, gas flow 12 (L/min), capillary voltage 3500 V. Data were processed using Agilent MassHunter Qualitative Analysis 10.0. Peaks were extracted by *m/z* within a 20 ppm error window. Metabolites were quantified by peak integration and area under the curve.

### Derivatization reactions with compound **5**

Derivatization reagent solutions were prepared immediately prior to each experiment by dissolving 4 µmol of 4,5-methylenedioxy-1,2-phenylenediamine dihydrochloride (Sigma Aldrich) and 56 µmol of sodium dithionite (Fisher Chemical) into 4 mL of deionized water to yield a final stock solution of 1 mM compound **5**. This derivatization reagent solution was diluted to desired concentrations in control solution (14 mM sodium dithionite in water). To perform derivatization reactions, the following mixtures were prepared in screw capped glass vials: 1 mL of derivatization reagent solution and 1 mL of compound **1** (Jubilant Pharma) or 1 mL of derivatization reagent solution and 1 mL of compound **4** (AK Scientific), each diluted to the concentrations designated in figure legends. For mammalian AI-2 mimic derivatization reactions, 50 µL of samples prepared as above were combined with 50 µL of a 7 mM solution of compound **5**. In all cases, mixtures were incubated at 55°C for 40 min. Control reactions were prepared identically except that the derivatization reagent solution was replaced by 1 mL of deionized water. Reactions were terminated by cooling on ice for 5 min. To measure the quinoxaline product, aliquots of reactions were placed into wells of black-sided, clear-bottom 96-well plates (Corning) and fluorescence intensity was measured in a Synergy plate-reader (BioTek) (excitation/emission, 355 nm/393 nm).

### Reactivity guided metabolomics

Derivatization reagent solutions were prepared immediately prior to each experiment by dissolving 12.6 µmol of 4,5-methylenedioxy-1,2-phenylenediamine dihydrochloride (Sigma Aldrich) and 25.2 µmol of sodium dithionite (Fisher Chemical) into 1.8 mL of deionized water to yield a final stock solution of 7 mM compound **5**. To perform derivatization reactions, screw capped glass vials containing 100 µL of compound **5** solution and 100 µL of mammalian AI-2 mimic preparations or 100 µL of 1X PBS were incubated at 55°C for 40 min, cooled in ice for 5 min, and subjected to centrifugation at 13,000 rpm for 5 min to remove insoluble material. High resolution LC-MS analyses were performed using the parameters described above. Following LC-MS acquisition, Agilent .d files were converted into .mzXML files using MSCovertGUI (ProteoWizard). Data in the converted files were analyzed using XCMS Online (Scripps University) in pairwise comparisons using the UPLC/UHD Q-TOF parameters. The peak picking algorithm was centWave, with significant features identified using unpaired parametric *t*-test (Welch t-test) and a *p*-value threshold of 0.05. To identify highly significant molecular features, we sorted and filtered the data using the calculated *q* value, which provides the false discovery rate (FDR) in multiple hypothesis testing.

### NMR methods

Nuclear magnetic resonance (NMR) spectra were acquired at the Princeton University Department of Chemistry Facilities. ^1^H, ^13^C and ^11^B NMR spectra were collected in the triple resonance cryoprobe of a Bruker Avance III 500 MHz NMR spectrometer, and were calibrated using residual undeuterated solvent as an internal reference (H_2_O: ^1^H NMR = 4.79; CDCl_3_: ^1^H NMR = 7.26, ^13^C NMR = 77.16; acetone-D_6_: ^1^H NMR = 2.05, ^13^C NMR = 29.84; CD_3_OD: ^1^H NMR = 3.31, ^13^C NMR = 49.00). ^1^H NMR spectra were tabulated as follows: chemical shift, multiplicity (s = singlet, d = doublet, t = triplet, q = quartet, p = pentet, dd = doublet of doublets, dt = doublet of triplets, m = multiplet, br = broad), coupling constant (Hz), and number of protons. ^13^C NMR spectra were tabulated by observed peak, and no special nomenclature is used for equivalent carbons. All NMR data were analyzed with MestReNova software.

### LC-MS Quantitation of compounds **8** and **18** in mammalian AI-2 mimic samples

Mammalian AI-2 mimic samples were prepared by splitting cells into seeding densities of 2,000,000 cells/mL and incubated in 5 mL of 1X PBS for 48 h followed by harvest. For LC-MS quantitation of compound **8**, derivatization reactions were prepared using 100 µL of AI-2 mammalian mimic preparations mixed with 100 µL of a 7 mM solution of compound **5**. The reactions were incubated at 55°C for 40 min, cooled on ice for 5 min, and stored at -20°C until LC-MS analysis. A standard curve was prepared by mixing 100 µL solutions (100 nM-1 mM) of compound **8** with a 7 mM solution of compound **5**. For quantitation of compound **18**, 400 µL of AI-2 mammalian mimic preparations were mixed with a 200 µL solution of 0.1 M N,N-diphenylhydrazine and 400 µL of MeOH. The N,N-diphenylhydrazine solution was prepared immediately prior to use by dissolving 1 mmol of diphenylhydrazine hydrochloride and 3 mmol of triethylamine in 10 mL water/MeCN (1:1, v/v). N,N-diphenylhydrazine derivatization reactions were allowed to stand for 24 h at room temperature prior to LC-MS analysis. Subsequently, samples were dried *in vacuo* (Genevac HT6 S3i Evaporator) followed by resuspension in 100 µL of MeOH for LC-MS analysis. *N*,*N*-diphenylhydrazine was chosen as the derivatizing agent to transform the analyte into a xylulose-hydrazone for optimized LC-MS detection in positive mode. Concentrations of compound **18** were estimated from a standard curve generated from derivatizations of 400 µL compound **18** solutions (50 µM-10 mM) with a 0.1 M *N*,*N*-diphenylhydrazine solution.

## Supporting Information

The Supporting Information is available free of charge on the ACS Publications website. Supporting information (PDF) includes Figures S1-S17, Table S1-S3, detailed synthetic methods, and all NMR characterization data.

## Author Contributions

E.E.S., J.S.V., V.Y.Y., and J.Z.H. conducted experiments. All authors designed research and analyzed data. All authors contributed to manuscript preparation.

## Acknowledgments

We thank members of the Bassler laboratory for helpful discussions. The authors acknowledge financial support from the Howard Hughes Medical Institute (B.L.B.), the National Institutes of Health grant R37GM065859 (B.L.B.), the National Science Foundation grant MCB-2508324 (B.L.B.), the National Institutes of Health grant R21AT013123 (M.R.S.), and a Life Sciences Research Foundation Postdoctoral Fellowship funded by the Open Philanthropy Project (J.Z.H.). The authors declare that they have no competing financial interests. Table of Contents Graphic created using BioRender.

## For Table of Contents Only

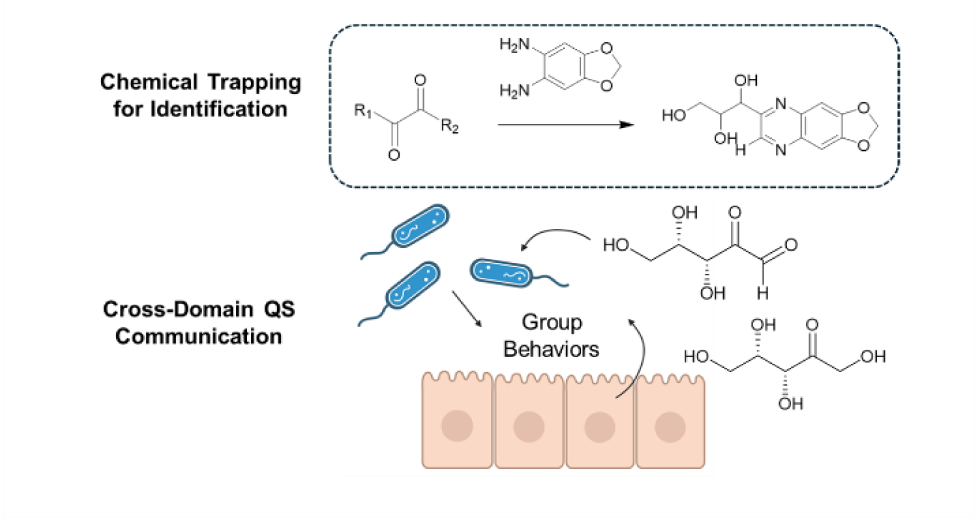

## Notes

### Competing Interest Statement

The authors have declared no competing interest.

## References

1. Fan, Y.; Pedersen, O., Gut Microbiota in Human Metabolic Health and Disease. Nat Rev Microbiol 2021, 19, 55–71.

2. Wirbel, J.; Pyl, P. T.; Kartal, E.; Zych, K.; Kashani, A.; Milanese, A.; Fleck, J. S.; Voigt, A. Y.; Palleja, A.; Ponnudurai, R.; Sunagawa, S.; Coelho, L. P.; Schrotz-King, P.; Vogtmann, E.; Habermann, N.; Niméus, E.; Thomas, A. M.; Manghi, P.; Gandini, S.; Serrano, D.; Mizutani, S.; Shiroma, H.; Shiba, S.; Shibata, T.; Yachida, S.; Yamada, T.; Waldron, L.; Naccarati, A.; Segata, N.; Sinha, R.; Ulrich, C. M.; Brenner, H.; Arumugam, M.; Bork, P.; Zeller, G., Meta-Analysis of Fecal Metagenomes Reveals Global Microbial Signatures That Are Specific for Colorectal Cancer. Nat. Med. 2019, 25, 679–689.

3. Lloyd-Price, J.; Arze, C.; Ananthakrishnan, A. N.; Schirmer, M.; Avila-Pacheco, J.; Poon, T. W.; Andrews, E.; Ajami, N. J.; Bonham, K. S.; Brislawn, C. J.; Casero, D.; Courtney, H.; Gonzalez, A.; Graeber, T. G.; Hall, A. B.; Lake, K.; Landers, C. J.; Mallick, H.; Plichta, D. R.; Prasad, M.; Rahnavard, G.; Sauk, J.; Shungin, D.; Vázquez-Baeza, Y.; White, R. A.; Bishai, J.; Bullock, K.; Deik, A.; Dennis, C.; Kaplan, J. L.; Khalili, H.; McIver, L. J.; Moran, C. J.; Nguyen, L.; Pierce, K. A.; Schwager, R.; Sirota-Madi, A.; Stevens, B. W.; Tan, W.; ten Hoeve, J. J.; Weingart, G.; Wilson, R. G.; Yajnik, V.; Braun, J.; Denson, L. A.; Jansson, J. K.; Knight, R.; Kugathasan, S.; McGovern, D. P. B.; Petrosino, J. F.; Stappenbeck, T. S.; Winter, H. S.; Clish, C. B.; Franzosa, E. A.; Vlamakis, H.; Xavier, R. J.; Huttenhower, C.; Investigators, I., Multi-omics of the Gut Microbial Ecosystem in Inflammatory Bowel Diseases. Nature 2019, 569, 655–662.

4. Zipperer, A.; Konnerth, M. C.; Laux, C.; Berscheid, A.; Janek, D.; Weidenmaier, C.; Burian, M.; Schilling, N. A.; Slavetinsky, C.; Marschal, M.; Willmann, M.; Kalbacher, H.; Schittek, B.; Brötz-Oesterhelt, H.; Grond, S.; Peschel, A.; Krismer, B., Human Commensals Producing a Novel Antibiotic Impair Pathogen Colonization. Nature 2016, 535, 511–6.

5. Torres Salazar, B. O.; Dema, T.; Schilling, N. A.; Janek, D.; Bornikoel, J.; Berscheid, A.; Elsherbini, A. M. A.; Krauss, S.; Jaag, S. J.; Lämmerhofer, M.; Li, M.; Alqahtani, N.; Horsburgh, M. J.; Weber, T.; Beltrán-Beleña, J. M.; Brötz-Oesterhelt, H.; Grond, S.; Krismer, B.; Peschel, A., Commensal Production of a Broad-Spectrum and Short-Lived Antimicrobial Peptide Polyene Eliminates Nasal Staphylococcus aureus. Nat. Microbiol. 2024, 9, 200–213.

6. Cohen, L. J.; Esterhazy, D.; Kim, S. H.; Lemetre, C.; Aguilar, R. R.; Gordon, E. A.; Pickard, A. J.; Cross, J. R.; Emiliano, A. B.; Han, S. M.; Chu, J.; Vila-Farres, X.; Kaplitt, J.; Rogoz, A.; Calle, P. Y.; Hunter, C.; Bitok, J. K.; Brady, S. F., Commensal Bacteria Make GPCR Ligands That Mimic Human Signalling Molecules. Nature 2017, 549, 48–53.

7. Cao, Y.; Oh, J.; Xue, M.; Huh, W. J.; Wang, J.; Gonzalez-Hernandez, J. A.; Rice, T. A.; Martin, A. L.; Song, D.; Crawford, J. M.; Herzon, S. B.; Palm, N. W., Commensal Microbiota From Patients with Inflammatory Bowel Disease Produce Genotoxic Metabolites. Science 2022, 378, eabm3233.

8. Lee, R.; Ptolemy, A. S.; Niewczas, L.; Britz-McKibbin, P., Integrative Metabolomics for Characterizing Unknown Low-Abundance Metabolites by Capillary Electrophoresis-Mass Spectrometry with Computer Simulations. Anal. Chem. 2007, 79, 403–415.

9. Xu, F.; Wu, Y.; Zhang, C.; Davis, K. M.; Moon, K.; Bushin, L. B.; Seyedsayamdost, M. R., A Genetics-Free Method for High-Throughput Discovery of Cryptic Microbial Metabolites. Nat. Chem. Biol. 2019, 15, 161–168.

10. Jiang, Y.; Stornetta, A.; Villalta, P. W.; Wilson, M. R.; Boudreau, P. D.; Zha, L.; Balbo, S.; Balskus, E. P., Reactivity of an Unusual Amidase May Explain Colibactin’s DNA Cross-Linking Activity. J. Am. Chem. Soc. 2019, 141, 11489–11496.

11. Xue, M.; Kim, C. S.; Healy, A. R.; Wernke, K. M.; Wang, Z.; Frischling, M. C.; Shine, E. E.; Wang, W.; Herzon, S. B.; Crawford, J. M., Structure Elucidation of Colibactin and its DNA Cross-links. Science 2019, 365.

12. Zhou, T.; Hirayama, Y.; Tsunematsu, Y.; Suzuki, N.; Tanaka, S.; Uchiyama, N.; Goda, Y.; Yoshikawa, Y.; Iwashita, Y.; Sato, M.; Miyoshi, N.; Mutoh, M.; Ishikawa, H.; Sugimura, H.; Wakabayashi, K.; Watanabe, K., Isolation of New Colibactin Metabolites from Wild-Type Escherichia coli and In Situ Trapping of a Mature Colibactin Derivative. J. Am. Chem. Soc. 2021, 143, 5526–5533.

13. Gentry, E. C.; Collins, S. L.; Panitchpakdi, M.; Belda-Ferre, P.; Stewart, A. K.; Carrillo Terrazas, M.; Lu, H.-h.; Zuffa, S.; Yan, T.; Avila-Pacheco, J.; Plichta, D. R.; Aron, A. T.; Wang, M.; Jarmusch, A. K.; Hao, F.; Syrkin-Nikolau, M.; Vlamakis, H.; Ananthakrishnan, A. N.; Boland, B. S.; Hemperly, A.; Vande Casteele, N.; Gonzalez, F. J.; Clish, C. B.; Xavier, R. J.; Chu, H.; Baker, E. S.; Patterson, A. D.; Knight, R.; Siegel, D.; Dorrestein, P. C., Reverse Metabolomics for the Discovery of Chemical Structures from Humans. Nature 2024, 626, 419–426.

14. Colby, S. M.; Nuñez, J. R.; Hodas, N. O.; Corley, C. D.; Renslow, R. R., Deep Learning to Generate in Silico Chemical Property Libraries and Candidate Molecules for Small Molecule Identification in Complex Samples. Anal. Chem. 2020, 92, 1720–1729.

15. Vargas, F.; Weldon, K. C.; Sikora, N.; Wang, M.; Zhang, Z.; Gentry, E. C.; Panitchpakdi, M. W.; Caraballo-Rodríguez, A. M.; Dorrestein, P. C.; Jarmusch, A. K., Protocol for Community-Created Public MS/MS Reference Spectra Within the Global Natural Products Social Molecular Networking Infrastructure. Rapid Commun. Mass Spectrom 2020, 34, e8725.

16. Covington, B. C.; Seyedsayamdost, M. R., Unlocking Hidden Treasures: the Evolution of High-Throughput Mass Spectrometry in Screening for Cryptic Natural Products. Nat. Prod. Rep. 2025, 42, 956–964.

17. Viant, M. R.; Kurland, I. J.; Jones, M. R.; Dunn, W. B., How Close Are We to Complete Annotation of Metabolomes? Curr. Opin. Chem. Biol. 2017, 36, 64–69.

18. Peisl, B. Y. L.; Schymanski, E. L.; Wilmes, P., Dark Matter in Host-Microbiome Metabolomics: Tackling the Unknowns–A Review. Anal. Chim. Acta 2018, 1037, 13–27.

19. Mukherjee, S.; Bassler, B. L., Bacterial Quorum Sensing in Complex and Dynamically Changing Environments. Nat. Rev. Microbiol. 2019, 17, 371–382.

20. Papenfort, K.; Bassler, B. L., Quorum Sensing Signal–Response Systems in Gram-Negative Bacteria. Nat. Rev. Microbiol. 2016, 14, 576–588.

21. Oliveira, R. A.; Cabral, V.; Torcato, I.; Xavier, K. B., Deciphering the Quorum-Sensing Lexicon of the Gut Microbiota. Cell Host Microbe 2023, 31, 500–512.

22. Schauder, S.; Shokat, K.; Surette, M. G.; Bassler, B. L., The LuxS Family of Bacterial Autoinducers: Biosynthesis Of a Novel Quorum-Sensing Signal Molecule. Mol. Microbiol. 2001, 41, 463–76.

23. Surette, M. G.; Miller, M. B.; Bassler, B. L., Quorum Sensing in Escherichia coli, Salmonella typhimurium, and Vibrio harveyi: a New Family of Genes Responsible for Autoinducer Production. Proc. Natl. Acad. Sci. U.S.A. 1999, 96, 1639–44.

24. Xavier, K. B.; Bassler, B. L., Interference with AI-2-Mediated Bacterial Cell–Cell Communication. Nature 2005, 437, 750–753.

25. Globisch, D.; Lowery, C. A.; McCague, K. C.; Janda, K. D., Uncharacterized 4,5-Dihydroxy-2,3-Pentanedione (DPD) Molecules Revealed Through NMR Spectroscopy: Implications for a Greater Signaling Diversity in Bacterial Species. Angew. Chem., Int. Ed. 2012, 51, 4204–4208.

26. Meijler, M. M.; Hom, L. G.; Kaufmann, G. F.; McKenzie, K. M.; Sun, C.; Moss, J. A.; Matsushita, M.; Janda, K. D., Synthesis and Biological Validation of a Ubiquitous Quorum-Sensing Molecule. Angew. Chem., Int. Ed. 2004, 43, 2106–8.

27. Miller, S. T.; Xavier, K. B.; Campagna, S. R.; Taga, M. E.; Semmelhack, M. F.; Bassler, B. L.; Hughson, F. M., Salmonella typhimurium Recognizes a Chemically Distinct Form of the Bacterial Quorum-Sensing Signal AI-2. Mol. Cell. 2004, 15, 677–87.

28. Semmelhack, M. F.; Campagna, S. R.; Federle, M. J.; Bassler, B. L., An Expeditious Synthesis of DPD and Boron Binding Studies. Org. Lett. 2005, 7, 569–572.

29. Chen, X.; Schauder, S.; Potier, N.; Van Dorsselaer, A.; Pelczer, I.; Bassler, B. L.; Hughson, F. M., Structural Identification of a Bacterial Quorum-Sensing Signal Containing Boron. Nature 2002, 415, 545–9.

30. Pereira Catarina, S.; de Regt Anna, K.; Brito Patrícia, H.; Miller Stephen, T.; Xavier Karina, B., Identification of Functional LsrB-Like Autoinducer-2 Receptors. J. Bacteriol. 2009, 191, 6975–6987.

31. Torcato, I. M.; Kasal, M. R.; Brito, P. H.; Miller, S. T.; Xavier, K. B., Identification of Novel Autoinducer-2 Receptors in Clostridia Reveals Plasticity in the Binding Site of the LsrB Receptor Family. J. Biol. Chem. 2019, 294, 4450–4463.

32. Zhang, L.; Li, S.; Liu, X.; Wang, Z.; Jiang, M.; Wang, R.; Xie, L.; Liu, Q.; Xie, X.; Shang, D.; Li, M.; Wei, Z.; Wang, Y.; Fan, C.; Luo, Z.-Q.; Shen, X., Sensing of Autoinducer-2 by Functionally Distinct Receptors in Prokaryotes. Nat. Commun. 2020, 11, 5371.

33. Fan, Q.; Sun, H.; Lin, X.; Yang, W.; Shen, X.; Zhang, L., Autoinducer-2-Mediated Communication Network within Human Gut Microbiota. ISME J 2025, 19.

34. Antunes, L. C.; Ferreira, L. Q.; Ferreira, E. O.; Miranda, K. R.; Avelar, K. E.; Domingues, R. M.; Ferreira, M. C., Bacteroides Species Produce Vibrio harveyi Autoinducer 2-Related Molecules. Anaerobe 2005, 11, 295–301.

35. Lukás, F.; Gorenc, G.; Kopecný, J., Detection of Possible AI-2-Mediated Quorum Sensing System in Commensal Intestinal Bacteria. Folia Microbiol (Praha) 2008, 53, 221–4.

36. Thompson, J. A.; Oliveira, R. A.; Djukovic, A.; Ubeda, C.; Xavier, K. B., Manipulation of the Quorum Sensing Signal AI-2 Affects the Antibiotic-Treated Gut Microbiota. Cell Rep. 2015, 10, 1861–71.

37. Valastyan, J. S.; Kraml, C. M.; Pelczer, I.; Ferrante, T.; Bassler, B. L., Saccharomyces cerevisiae Requires CFF1 To Produce 4-Hydroxy-5-Methylfuran-3(2H)-One, a Mimic of the Bacterial Quorum-Sensing Autoinducer AI-2. mBio 2021, 12.

38. Ismail, Anisa S.; Valastyan, Julie S.; Bassler, Bonnie L., A Host-Produced Autoinducer-2 Mimic Activates Bacterial Quorum Sensing. Cell Host Microbe 2016, 19, 470–480.

39. Campagna, S. R.; Gooding, J. R.; May, A. L., Direct Quantitation of the Quorum Sensing Signal, Autoinducer-2, in Clinically Relevant Samples by Liquid Chromatography−Tandem Mass Spectrometry. Anal. Chem. 2009, 81, 6374–6381.

40. Xu, F.; Song, X.; Cai, P.; Sheng, G.; Yu, H., Quantitative Determination of AI-2 Quorum-Sensing Signal of Bacteria Using High Performance Liquid Chromatography–Tandem Mass Spectrometry. J. Environ. Sci. (China) 2017, 52, 204–209.

41. Rodrigues, M. V.; Ferreira, A.; Ramirez-Montoya, M.; Oliveira, R. A.; Defaix, R.; Kis, P.; Cabral, V.; Bronze, M. R.; Xavier, K. B.; Ventura, M. R., Manipulation and Quantification of the Levels of Autoinducer-2 Quorum Sensing Signal in the Mouse Gut. Bioorg Chem. 2025, 157, 108274.

42. Thiel, V.; Vilchez, R.; Sztajer, H.; Wagner-Döbler, I.; Schulz, S., Identification, Quantification, and Determination of the Absolute Configuration of the Bacterial Quorum-Sensing Signal Autoinducer-2 by Gas Chromatography-Mass Spectrometry. ChemBioChem 2009, 10, 479–85.

43. Hara, S.; Yamaguchi, M.; Takemori, Y.; Yoshitake, T.; Nakamura, M., 1,2-Diamino-4,5-Methylenedioxybenzene as a Highly Sensitive Fluorogenic Reagent for α-Dicarbonyl Compounds. Anal. Chim. Acta 1988, 215, 267–276.

44. Long, T.; Tu, K. C.; Wang, Y.; Mehta, P.; Ong, N. P.; Bassler, B. L.; Wingreen, N. S., Quantifying the Integration of Quorum-Sensing Signals with Single-Cell Resolution. PLoS Biol. 2009, 7, e68.

45. Henning, C.; Liehr, K.; Girndt, M.; Ulrich, C.; Glomb, M. A., Extending the Spectrum of α-Dicarbonyl Compounds In Vivo. J. Biol. Chem. 2014, 289, 28676–88.

46. Lopez, R.; Fernandez-Mayoralas, A., Enzymic Beta-Galactosidation of Modified Monosaccharides: Study of the Enzyme Selectivity for the Acceptor and Its Application to the Synthesis of Disaccharides. J. Org. Chem. 1994, 59, 737–745.

47. Phanumartwiwath, A.; Hornsby, T. W.; Jamalis, J.; Bailey, C. D.; Willis, C. L., Silyl Migrations in D-Xylose Derivatives: Total Synthesis of a Marine Quinoline Alkaloid. Org. Lett. 2013, 15, 5734–5737.

48. Volc, J.; Sedmera, P.; Halada, P.; Přikrylová, V.; Haltrich, D., Double Oxidation of D-Xylose to D-Glycero-Pentos-2,3-Diulose (2,3-Diketo-D-Xylose) by Pyranose Dehydrogenase from the Mushroom Agaricusbisporus. Carbohydr. Res. 2000, 329, 219–225.

49. Lowery, C. A.; McKenzie, K. M.; Qi, L.; Meijler, M. M.; Janda, K. D., Quorum Sensing in Vibrio harveyi: Probing the Specificity of the LuxP Binding Site. Bioorg. Med. Chem. Lett. 2005, 15, 2395–8.

50. Miller, M. B.; Skorupski, K.; Lenz, D. H.; Taylor, R. K.; Bassler, B. L., Parallel Quorum Sensing Systems Converge to Regulate Virulence in Vibrio cholerae. Cell 2002, 110, 303–314.

51. Zhu, J.; Miller, M. B.; Vance, R. E.; Dziejman, M.; Bassler, B. L.; Mekalanos, J. J., Quorum-sensing regulators control virulence gene expression in Vibrio cholerae. Proc Natl Acad Sci U S A 2002, 99, 3129–34.

52. Singh, P. K.; Bartalomej, S.; Hartmann, R.; Jeckel, H.; Vidakovic, L.; Nadell, C. D.; Drescher, K., Vibrio cholerae Combines Individual and Collective Sensing to Trigger Biofilm Dispersal. Curr Biol 2017, 27, 3359–3366 e7.

53. Bridges, A. A.; Bassler, B. L., The intragenus and interspecies quorum-sensing autoinducers exert distinct control over Vibrio cholerae biofilm formation and dispersal. PLOS Biology 2019, 17, e3000429.

54. van den Berg, R.; Peters, J. A.; van Bekkum, H., The structure and (local) stability constants of borate esters of mono- and di-saccharides as studied by 11B and 13C NMR spectroscopy. Carbohydrate Research 1994, 253, 1–12.

55. Semmelhack, M. F.; Campagna, S. R.; Hwa, C.; Federle, M. J.; Bassler, B. L., Boron Binding with the Quorum Sensing Signal AI-2 and Analogues. Organic Letters 2004, 6, 2635–2637.

56. Smith, P. M.; Howitt, M. R.; Panikov, N.; Michaud, M.; Gallini, C. A.; Bohlooly, Y. M.; Glickman, J. N.; Garrett, W. S., The Microbial Metabolites, Short-Chain Fatty Acids, Regulate Colonic Treg Cell Homeostasis. Science 2013, 341, 569–73.

57. Paik, D.; Yao, L.; Zhang, Y.; Bae, S.; D’Agostino, G. D.; Zhang, M.; Kim, E.; Franzosa, E. A.; Avila-Pacheco, J.; Bisanz, J. E.; Rakowski, C. K.; Vlamakis, H.; Xavier, R. J.; Turnbaugh, P. J.; Longman, R. S.; Krout, M. R.; Clish, C. B.; Rastinejad, F.; Huttenhower, C.; Huh, J. R.; Devlin, A. S., Human Gut Bacteria Produce Τ(Η)17-Modulating Bile Acid Metabolites. Nature 2022, 603, 907–912.

58. Quinn, R. A.; Melnik, A. V.; Vrbanac, A.; Fu, T.; Patras, K. A.; Christy, M. P.; Bodai, Z.; Belda-Ferre, P.; Tripathi, A.; Chung, L. K.; Downes, M.; Welch, R. D.; Quinn, M.; Humphrey, G.; Panitchpakdi, M.; Weldon, K. C.; Aksenov, A.; da Silva, R.; Avila-Pacheco, J.; Clish, C.; Bae, S.; Mallick, H.; Franzosa, E. A.; Lloyd-Price, J.; Bussell, R.; Thron, T.; Nelson, A. T.; Wang, M.; Leszczynski, E.; Vargas, F.; Gauglitz, J. M.; Meehan, M. J.; Gentry, E.; Arthur, T. D.; Komor, A. C.; Poulsen, O.; Boland, B. S.; Chang, J. T.; Sandborn, W. J.; Lim, M.; Garg, N.; Lumeng, J. C.; Xavier, R. J.; Kazmierczak, B. I.; Jain, R.; Egan, M.; Rhee, K. E.; Ferguson, D.; Raffatellu, M.; Vlamakis, H.; Haddad, G. G.; Siegel, D.; Huttenhower, C.; Mazmanian, S. K.; Evans, R. M.; Nizet, V.; Knight, R.; Dorrestein, P. C., Global Chemical Effects of the Microbiome Include New Bile-Acid Conjugations. Nature 2020, 579, 123–129.

59. Dodd, D.; Spitzer, M. H.; Van Treuren, W.; Merrill, B. D.; Hryckowian, A. J.; Higginbottom, S. K.; Le, A.; Cowan, T. M.; Nolan, G. P.; Fischbach, M. A.; Sonnenburg, J. L., A Gut Bacterial Pathway Metabolizes Aromatic Amino Acids into Nine Circulating Metabolites. Nature 2017, 551, 648–652.

60. Zelante, T.; Iannitti, R. G.; Cunha, C.; De Luca, A.; Giovannini, G.; Pieraccini, G.; Zecchi, R.; D’Angelo, C.; Massi-Benedetti, C.; Fallarino, F.; Carvalho, A.; Puccetti, P.; Romani, L., Tryptophan Catabolites from Microbiota Engage Aryl Hydrocarbon Receptor and Balance Mucosal Reactivity via Interleukin-22. Immunity 2013, 39, 372–85.

61. Rooks, M. G.; Garrett, W. S., Gut Microbiota, Metabolites and Host Immunity. Nat. Rev. Immunol. 2016, 16, 341–52.

62. Hooper, L. V.; Littman, D. R.; Macpherson, A. J., Interactions Between the Microbiota and the Immune System. Science 2012, 336, 1268–1273.

63. Nemet, I.; Monnier, V. M., Vitamin C Degradation Products and Pathways in the Human Lens. J. Biol. Chem. 2011, 286, 37128–36.

64. Nagaraj, R. H.; Sell, D. R.; Prabhakaram, M.; Ortwerth, B. J.; Monnier, V. M., High Correlation Between Pentosidine Protein Crosslinks and Pigmentation Implicates Ascorbate Oxidation in Human Lens Senescence and Cataractogenesis. Proc. Natl. Acad. Sci. U.S.A. 1991, 88, 10257–61.

65. Reihl, O.; Lederer, M. O.; Schwack, W., Characterization and Detection of Lysine–Arginine Cross-Links Derived from Dehydroascorbic Acid. Carbohydr. Res. 2004, 339, 483–491.

66. Whiting, G. C.; Coggins, R. A., Formation of L-xylosone From Ascorbic Acid. Nature 1960, 185, 843–4.

67. Shin, D. B.; Feather, M. S., 3-Deoxy-L-Glycero-Pentos-2-ulose (3-Deoxy-L-Xylosone) and L-Threo-Pentos-2-ulose (L-Xylosone) as Intermediates in the Degradation of L-Ascorbic Acid. Carbohydr. Res. 1990, 208, 246–250.

68. Giffhorn, F., Fungal Pyranose Oxidases: Occurrence, Properties and Biotechnical Applications in Carbohydrate Chemistry. Appl. Microbiol. Biotechnol. 2000, 54, 727–40.

69. Abrera, A. T.; Sützl, L.; Haltrich, D., Pyranose Oxidase: A Versatile Sugar Oxidoreductase for Bioelectrochemical Applications. Bioelectrochemistry 2020, 132, 107409.

70. Sützl, L.; Foley, G.; Gillam, E. M. J.; Bodén, M.; Haltrich, D., The GMC Superfamily of Oxidoreductases Revisited: Analysis and Evolution of Fungal GMC Oxidoreductases. Biotechnol Biofuels 2019, 12, 118.

71. Santema, L. L.; Rozeboom, H. J.; Borger, V. P.; Kaya, S. G.; Fraaije, M. W., Identification of a Robust Bacterial Pyranose Oxidase that Displays an Unusual pH Dependence. J. Biol. Chem. 2024, 300, 107885.

72. Sochor, M.; Baquer, N. Z.; McLean, P., Glucose Overutilization in Diabetes: Evidence from Studies on the Changes in Hexokinase, the Pentose Phosphate Pathway and Glucuronate-Xylulose Pathway in Rat Kidney Cortex in Diabetes. Biochem Biophys Res Commun. 1979, 86, 32–9.

73. Malatesta, M.; De Rito, C.; Gasparini, F.; Merici, G.; Dell’Accantera, D.; Quilici, G.; Sansone, F.; Percudani, R., C11orf54 Catalyzes L-Xylulose Formation in Human Metabolism. Proc. Natl. Acad. Sci. U.S.A. 2025, 122, e2506597122.

74. Pierce, S. B.; Spurrell, C. H.; Mandell, J. B.; Lee, M. K.; Zeligson, S.; Bereman, M. S.; Stray, S. M.; Fokstuen, S.; MacCoss, M. J.; Levy-Lahad, E.; King, M. C.; Motulsky, A. G., Garrod’s Fourth Inborn Error of Metabolism Solved by the Identification of Mutations Causing Pentosuria. Proc. Natl. Acad. Sci. U.S.A. 2011, 108, 18313–7.

75. Odani, H.; Asami, J.; Ishii, A.; Oide, K.; Sudo, T.; Nakamura, A.; Miyata, N.; Otsuka, N.; Maeda, K.; Nakagawa, J., Suppression of Renal Alpha-Dicarbonyl Compounds Generated Following Ureteral Obstruction by Kidney-Specific Alpha-Dicarbonyl/L-Xylulose Reductase. Ann N Y Acad Sci 2008, 1126, 320–4.

76. Asami, J.; Odani, H.; Ishii, A.; Oide, K.; Sudo, T.; Nakamura, A.; Miyata, N.; Otsuka, N.; Maeda, K.; Nakagawa, J., Suppression of AGE Precursor Formation Following Unilateral Ureteral Obstruction in Mouse Kidneys by Transgenic Expression of Alpha-Dicarbonyl/L-Xylulose Reductase. Biosci Biotechnol Biochem 2006, 70, 2899–905.

77. Hubbs, A. F.; Fluharty, K. L.; Edwards, R. J.; Barnabei, J. L.; Grantham, J. T.; Palmer, S. M.; Kelly, F.; Sargent, L. M.; Reynolds, S. H.; Mercer, R. R.; Goravanahally, M. P.; Kashon, M. L.; Honaker, J. C.; Jackson, M. C.; Cumpston, A. M.; Goldsmith, W. T.; McKinney, W.; Fedan, J. S.; Battelli, L. A.; Munro, T.; Bucklew-Moyers, W.; McKinstry, K.; Schwegler-Berry, D.; Friend, S.; Knepp, A. K.; Smith, S. L.; Sriram, K., Accumulation of Ubiquitin and Sequestosome-1 Implicate Protein Damage in Diacetyl-Induced Cytotoxicity. Am J Pathol 2016, 186, 2887–2908.

78. Cox, C. L.; Tietz, J. I.; Sokolowski, K.; Melby, J. O.; Doroghazi, J. R.; Mitchell, D. A., Nucleophilic 1,4-Additions for Natural Product Discovery. ACS Chem. Biol. 2014, 9, 2014–2022.

79. Harris, L. A.; Mitchell, D. A., Reactivity-Based Screening for Natural Product Discovery. Methods Enzymol 2022, 665, 177–208.

80. Huang, Y. B.; Cai, W.; Del Rio Flores, A.; Twigg, F. F.; Zhang, W., Facile Discovery and Quantification of Isonitrile Natural Products via Tetrazine-Based Click Reactions. Anal. Chem. 2020, 92, 599–602.

81. Castro-Falcón, G.; Hahn, D.; Reimer, D.; Hughes, C. C., Thiol Probes To Detect Electrophilic Natural Products Based on Their Mechanism of Action. ACS Chem. Biol. 2016, 11, 2328–36.

82. Pfeifer, K.; Van Cura, D.; Wu, K. J. Y.; Balskus, E. P., Chemical Capture of Diazo Metabolites Reveals Biosynthetic Hydrazone Oxidation. bioRxiv 2025, 2025.05.24.655970.

83. Mittelmaier, S.; Fünfrocken, M.; Fenn, D.; Berlich, R.; Pischetsrieder, M., Quantification of the Six Major α-Dicarbonyl Contaminants in Peritoneal Dialysis Fluids by UHPLC/DAD/MSMS. Anal. Bioanal. Chem. 2011, 401, 1183–93.

84. Thornalley, P. J.; Langborg, A.; Minhas, H. S., Formation of Glyoxal, Methylglyoxal and 3-Deoxyglucosone in the Glycation of Proteins by Glucose. Biochem J. 1999, 344 *Pt* *1*, 109–16.

85. Cha, J.; Debnath, T.; Lee, K.-G., Analysis of α-Dicarbonyl Compounds and Volatiles Formed in Maillard Reaction Model Systems. Sci. Rep. 2019, 9, 5325.

