## Supplemental Information for "Discovery of a Human Metabolite that Mimics the Bacterial Quorum-Sensing Autoinducer AI-2"

### Table of Contents

Figures S1 to S17……………………………………………………………………..S2

Tables S1 to S3 ………………………………………………………………………S17

Synthetic methods and characterization data for all synthetic compounds…………………..S19

Supplementary References…………………………………………………………...S43

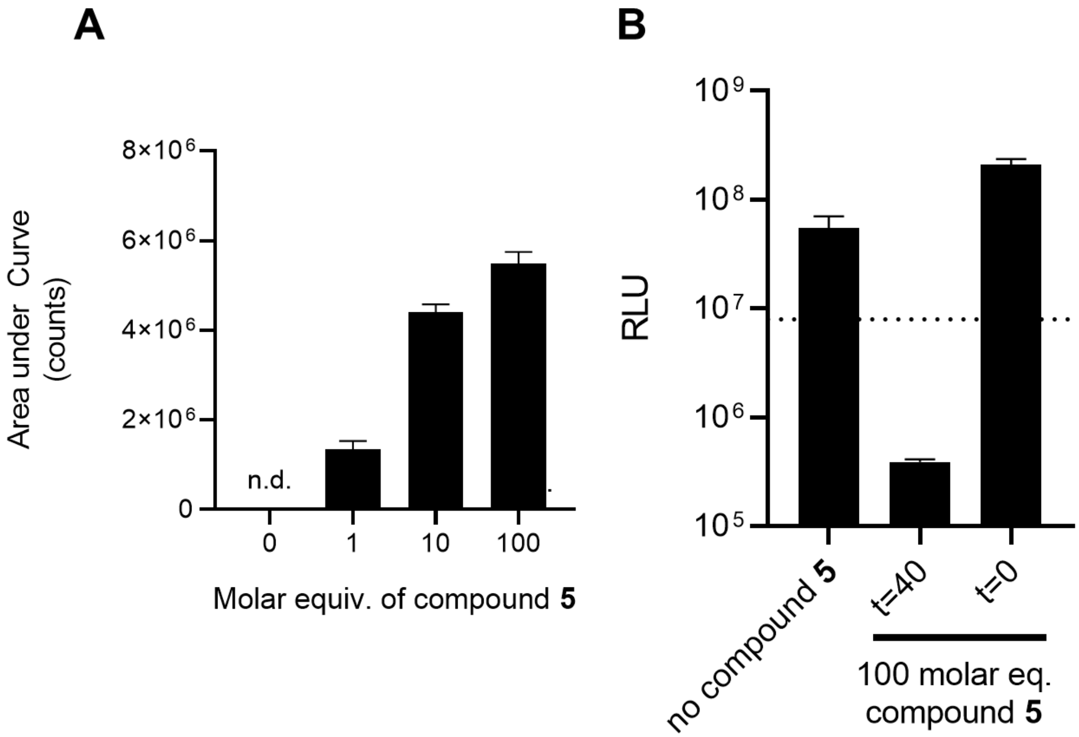


**Figure S1.** Optimization of DPD (**1**) derivatization with DMB (**5**). A) Relative quantitation of DPD-quinoxaline (**6**) measured by LC-MS. In each reaction, 5 nmol **1** was incubated with the indicated molar equivalents of **5**. Reactions were carried out at 55°C for 40 min. Data were quantified as area under the peaks for extracted *m/z* 249.1. Error bars denote standard deviations for biological replicates, $n = 3$. B) Bioluminescence output from the *V. harveyi* TL-26 bioassay strain. In each reaction, 5 nmol **1** was incubated alone or with 100 molar equivalents of **5**. t = 0 indicates that compound **5** and compound **1** were added simultaneously to the reporter strain without prior reaction. t = 40 min indicates that compound **5** and compound **1** were reacted at 55°C for 40 min before exposure to the reporter strain. In B, RLU denotes relative light units, which are bioluminescence/$OD_{600}$. n.d. = not detected.

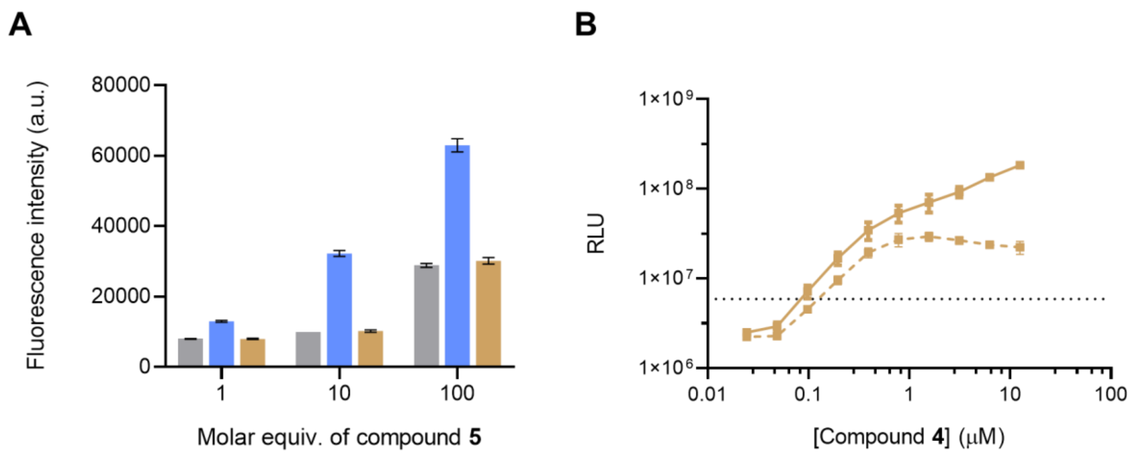


**Figure S2.** MHF, **4**, which lacks an a-diketone, is not derivatized by DMB, **5**. A) Fluorescence intensities of reaction mixtures containing water (gray), 10 nmol of **1** (blue) or 100 nmol of **4** (gold) incubated with the indicated molar equivalents of **5**. Production of a fluorescent quinoxaline product was detected at excitation/emission 355 nm/393 nm. B) Reactions containing 500 µM of **4** were incubated with either control solution (solid line) or 3 mM **5** (dashed line) for 40 min at 55°C followed by application in serial dilution to the *V. harveyi* TL-26 bioassay strain. RLU denotes relative light units, which are bioluminescence/$OD_{600}$, and the dotted line indicates the baseline light output in the bioassay. In both panels, error bars denote standard deviations for biological replicates, $n$ = 3.

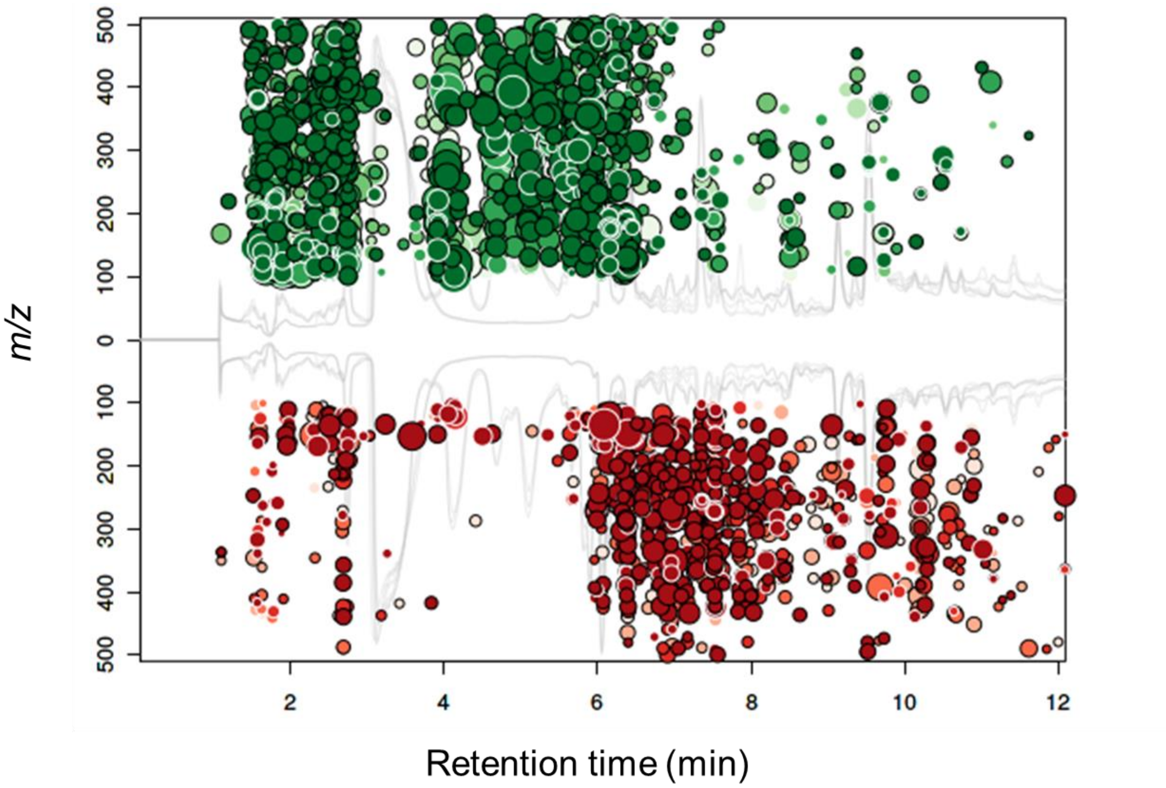


**Figure S3.** Metabolome cloud plot generated from XC-MS Online comparing mammalian AI-2 mimic samples derivatized with **5** versus PBS treated with **5**. Each point represents a distinct molecular feature, with point sizes proportional to fold-changes and colors corresponding to *p*-values ($n$ = 3 biological replicates). Darker colors indicate lower *p*-values. Retention times are represented by positions on the x-axis. Mass-to-charge ratios (*m/z*) are represented by positions on the y-axis. Points in green represent molecular features enriched in the AI-2 mimic sample treated with compound **5**. Points in red represent molecular features enriched in the control.

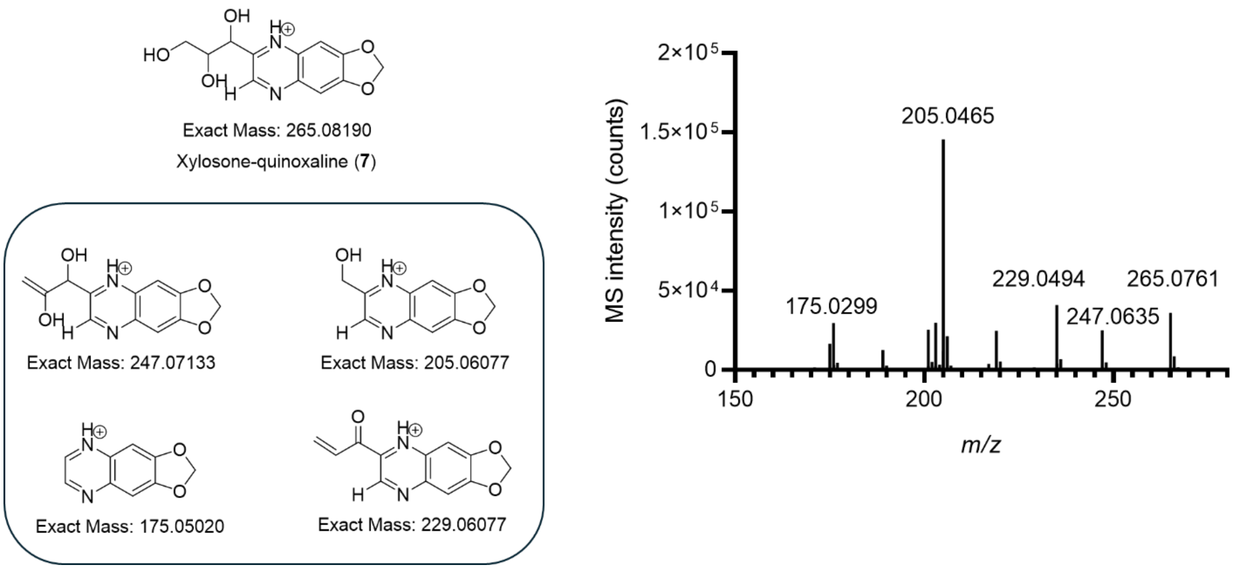


**Figure S4.** MS/MS spectrum of **7** resulting from synthesized **8** derivatized with **5**. Shown in the box are predicted fragment ion structures are shown with computed exact masses calculated for [M+H]. Observed *m/z* fragments are labeled in the spectrum.

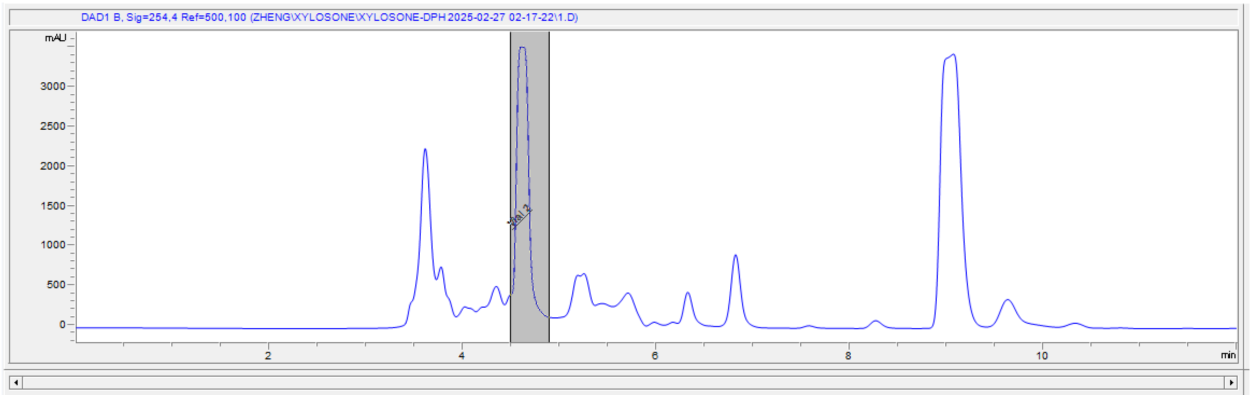


**Figure S5.** HPLC trace for the purification of L-xylosone DPH derivative.

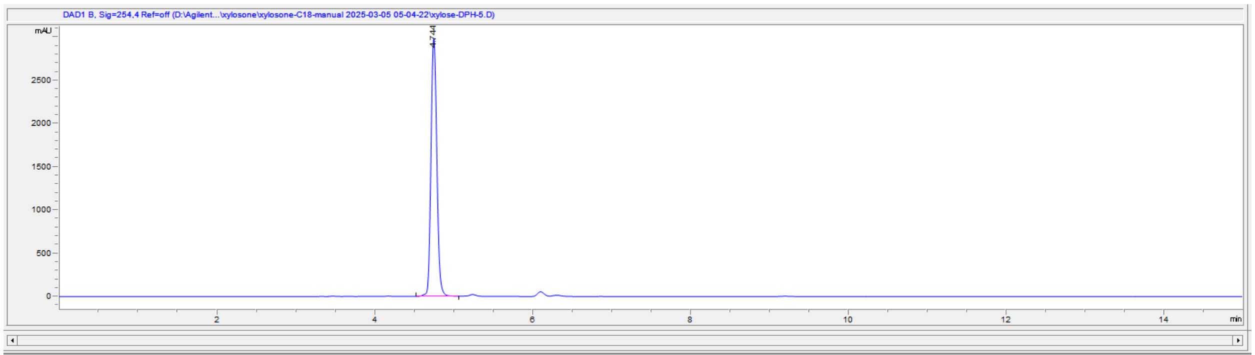


| File Information | |
|---|---|
| LC-File | xylose-DPH-5.D |
| File Path | D:\Agilent\Zheng\xylosone\xylosone-C18-manual 2025-03-05 05-0 |
| Date | 05-Mar-25, 05:38:19 |
| Sample | xylose-DPH-5 |
| Sample Info | |
| Barcode | |
| Operator | SYSTEM |
| Method | xylosone.M |
| Reference | |
| Analysis Time | 14.993 min |
| Sampling Rate | 0.0067 min (0.402 sec), 2250 datapoints |

| # | Time | Type | Area | Height | Width | Area% | Symmetry |
|---|---|---|---|---|---|---|---|
| 1 | 4.744 | BB | 15393 | 2985.2 | 0.0802 | 100.000 | 0.864 |

**Figure S6.** Representative HPLC trace for the 5 mg/mL stock of L-xylosone DPH derivative.

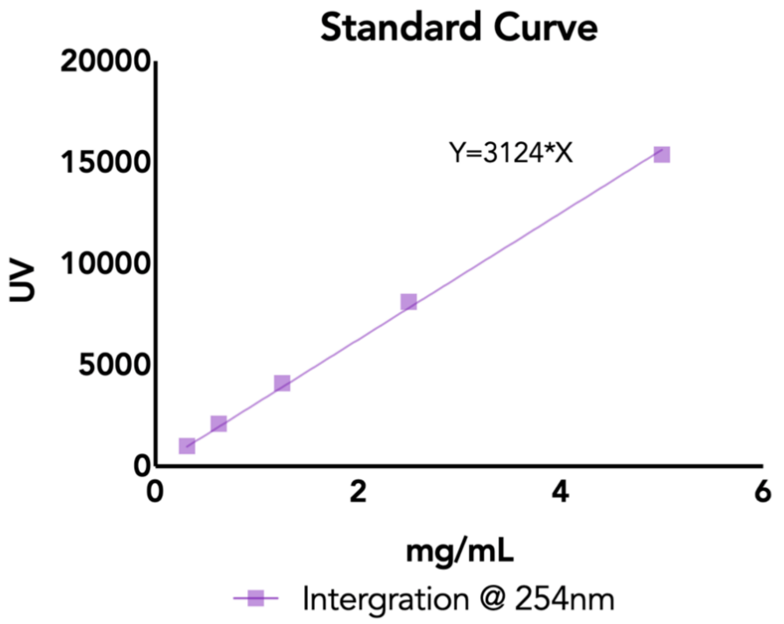


**Figure S7.** Standard curve for the HPLC-UV quantification of DPH-modified xylosone.

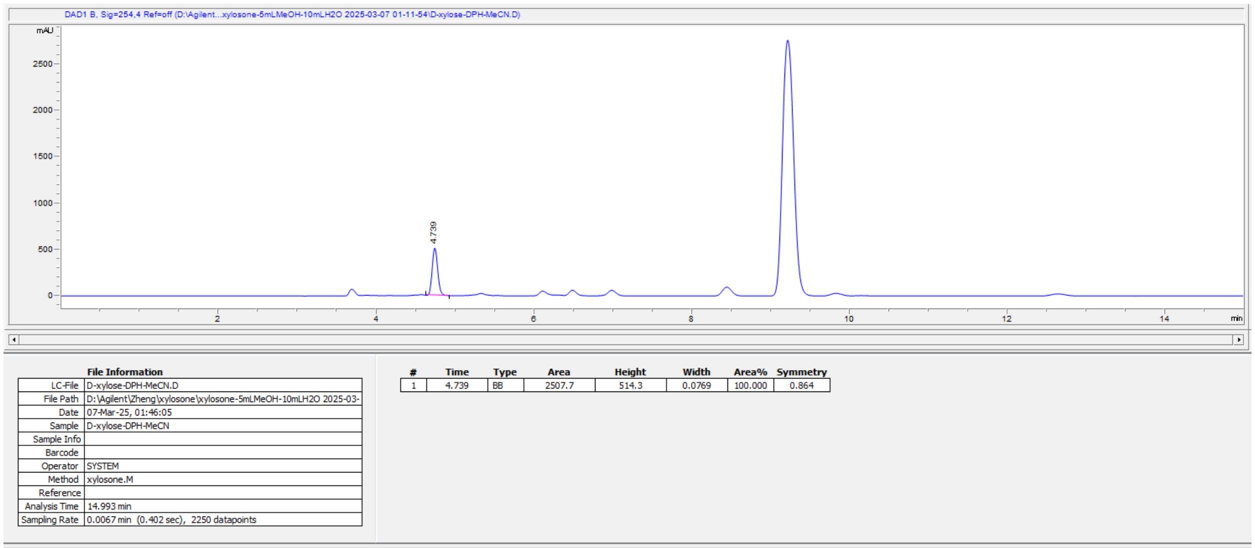


**Figure S8.** HPLC trace for DPH-modified D-xylosone. The concentration of the initial D-xylosone stock is calculated to be 6.4 mM.

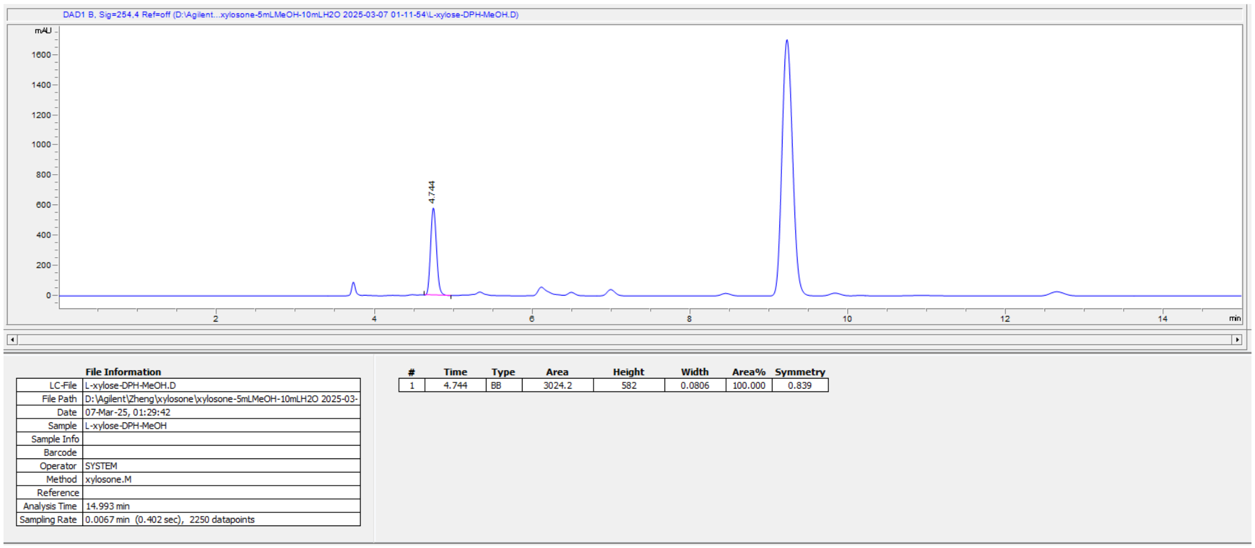


**Figure S9.** HPLC trace for DPH-modified L-xylosone. The concentration of the initial L-xylosone stock is calculated to be 7.7 mM.

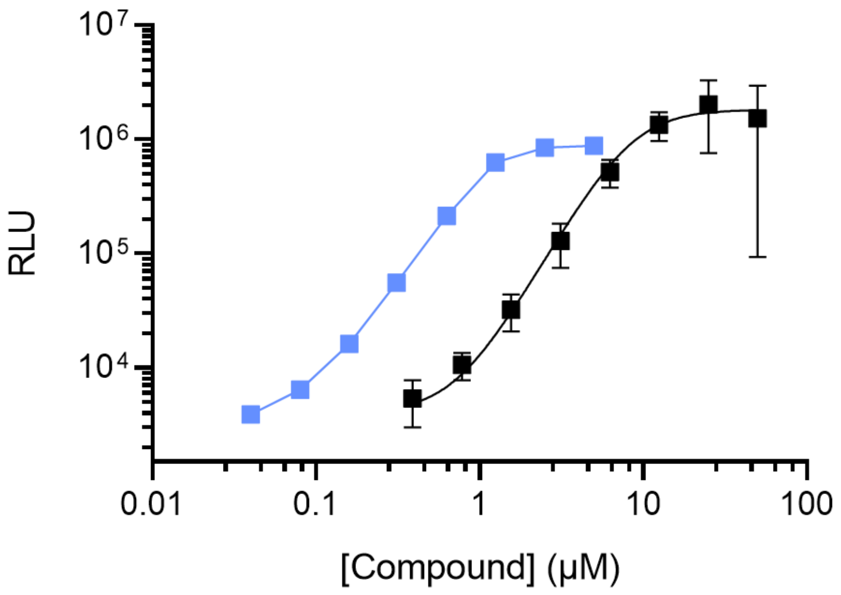


| | EC50 (μM) |
|---|---|
| Compound **1** | 1.0 |
| Compound **8** | 8.3 |

**Figure S10**. Light output from the *V. cholerae* AI-2 reporter in response to indicated amounts of compound **1** (blue) and **8** (black). The calculated EC50 values are shown in the table below. Error bars represent standard deviations of technical replicates, n = 3.

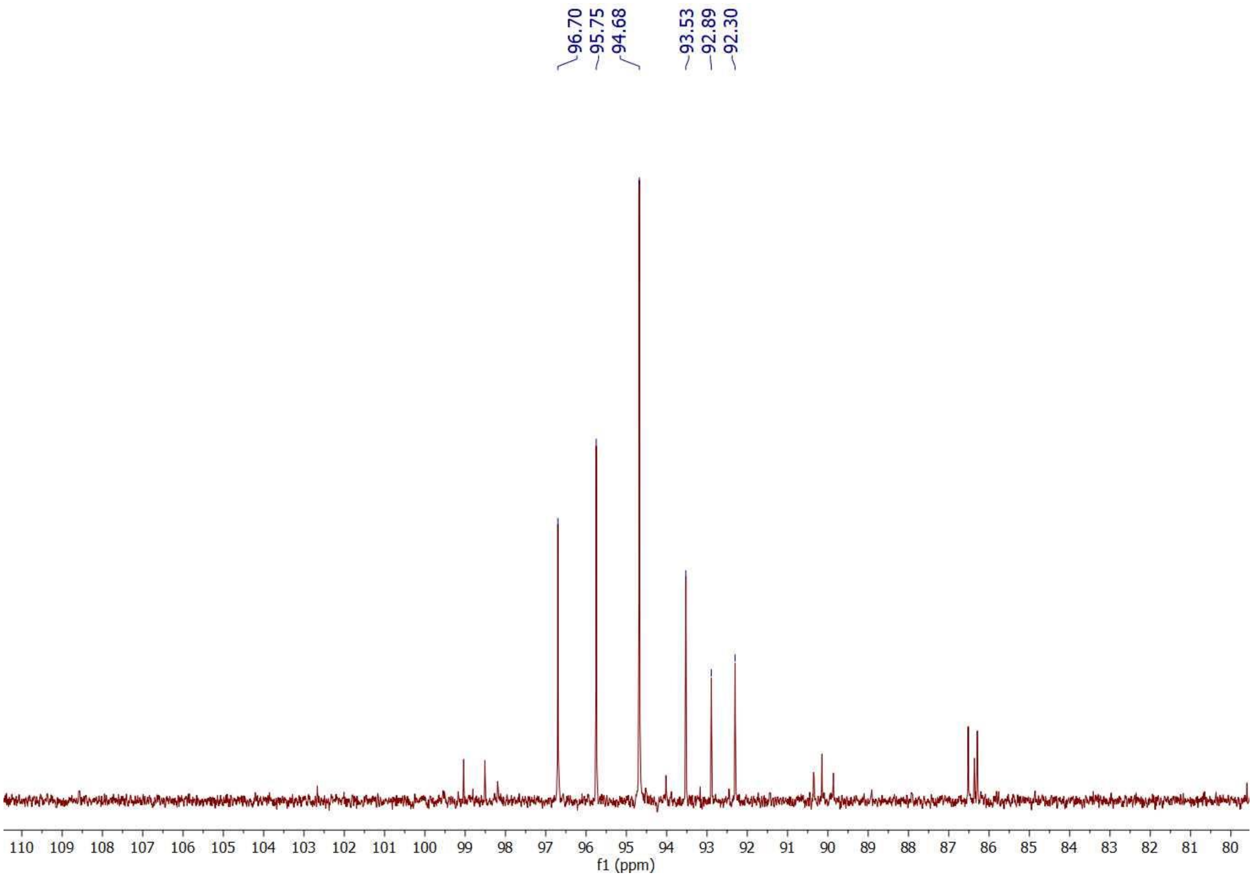


**Figure S11**. Expansion of the acetal region for $^{13}C$ NMR spectrum of compound **8**. Six major peaks are annotated with their corresponding chemical shifts in ppm. The spectrum was recorded in 95/5 $H_2O/D_2O$ mixture.

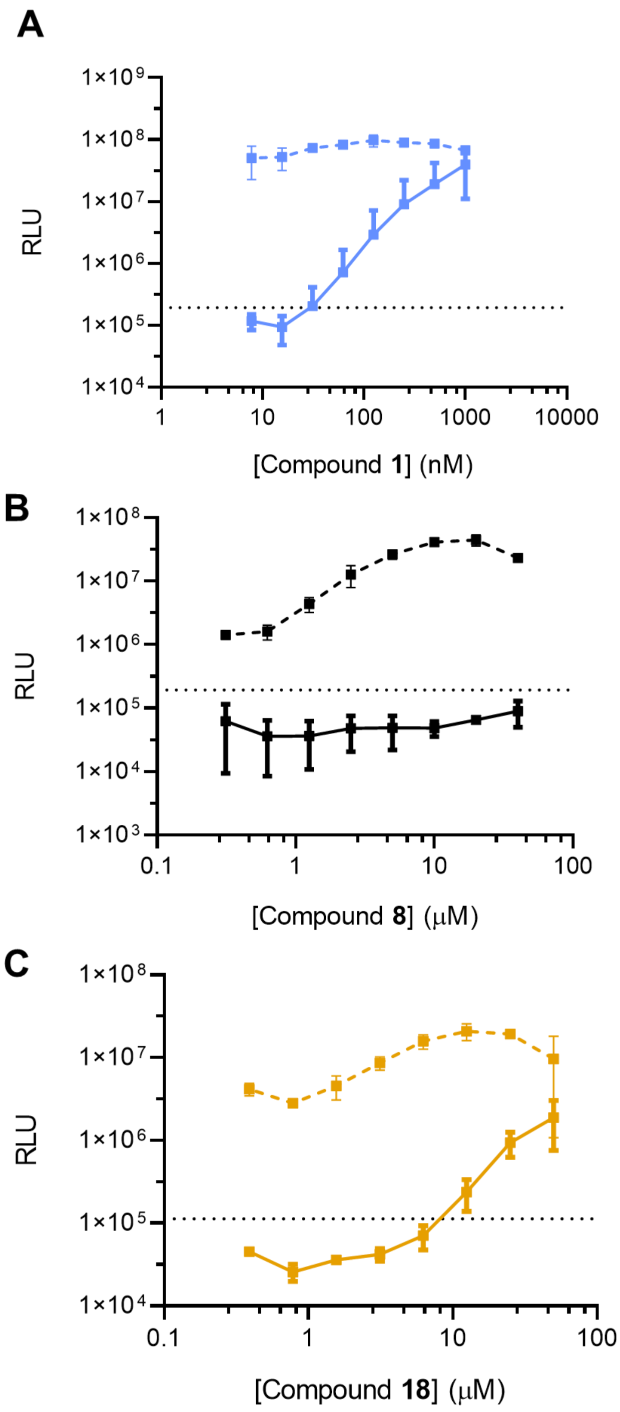


**Figure S12**. Bioluminescence output from the *V. harveyi* TL-26 bioassay strain in response to compounds **1** (A), **8** (B), and **18** (C). Solid and dashed lines indicate no addition or addition of 0.1 mM boric acid to the medium. Error bars denote standard deviations for biological replicates, *n* = 3.

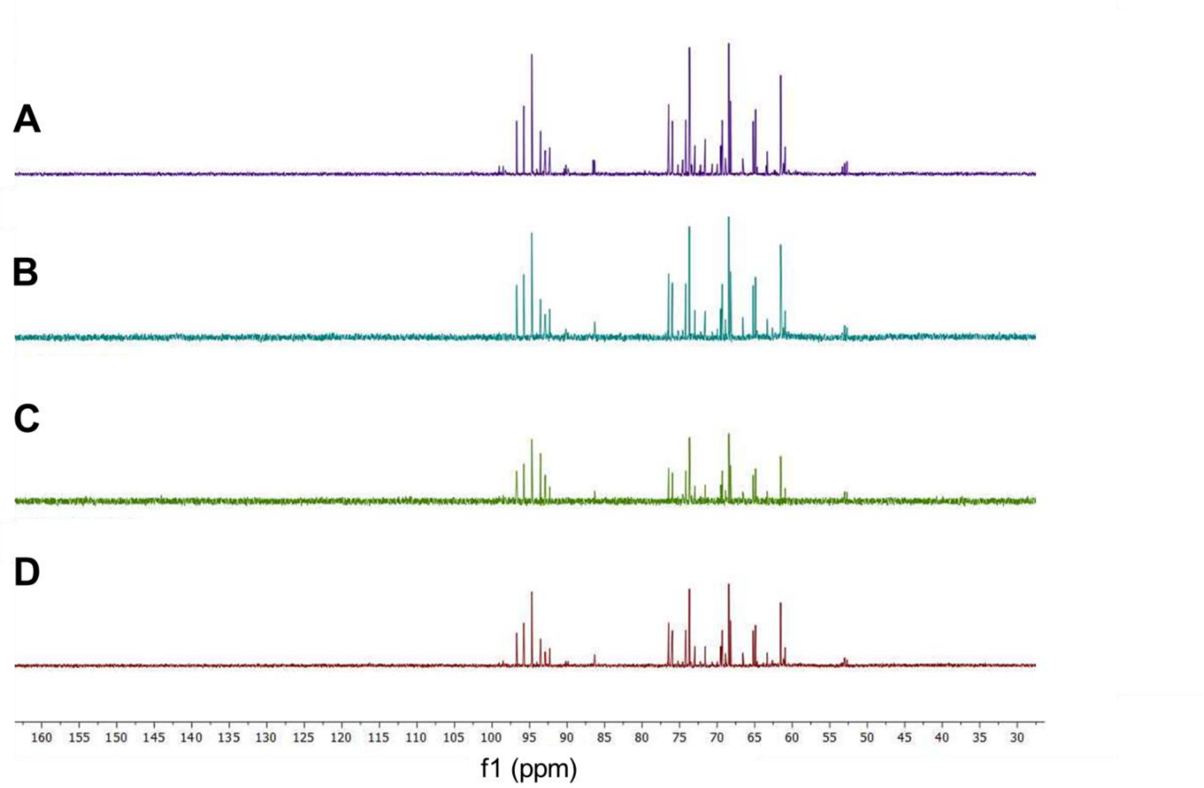


**Figure S13**. [13]C NMR comparison of 50 mM compound **8** titrated with different concentrations of boric acid. A) Compound **8** alone, B) 2:1 boric acid/compound **8**, C) 1:2 boric acid/compound **8**, D) 1:1 boric acid/**8**. All spectra were recorded in 95/5 $H_2O/D_2O$ mixture.

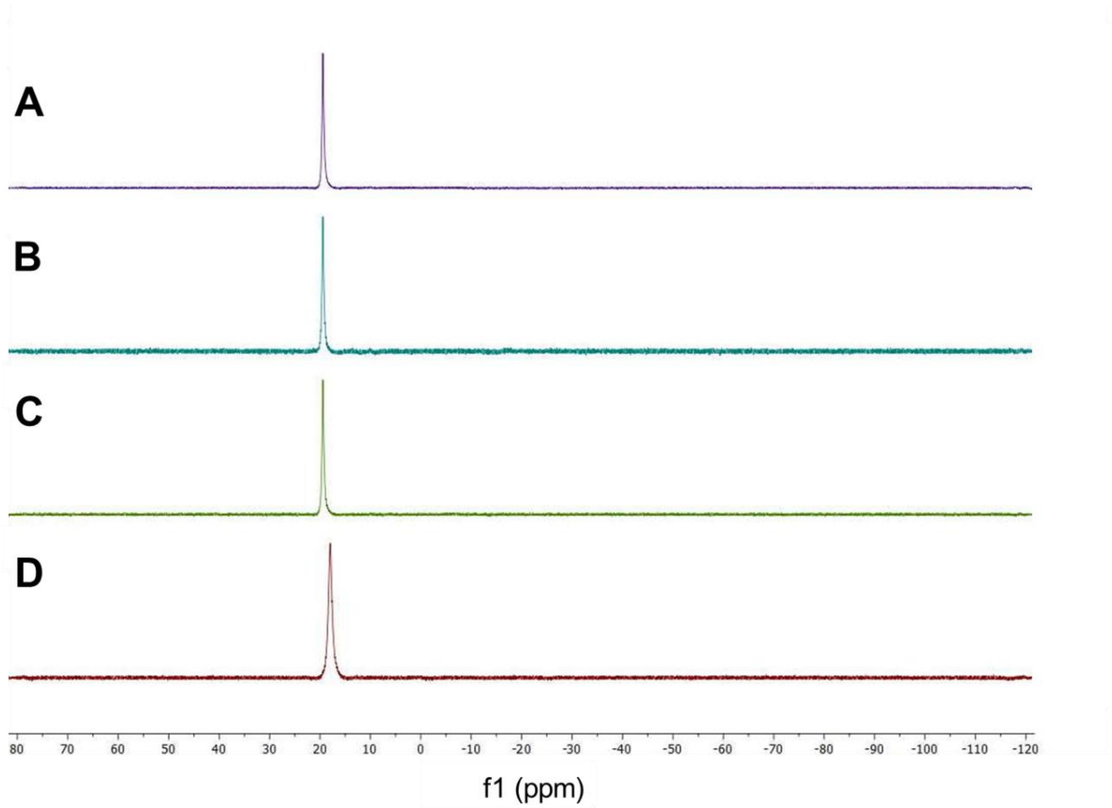


**Figure S14**. $^{11}$B NMR comparison of 50 mM compound **8** titrated with different concentrations of boric acid. A) 2:1 boric acid/compound **8**, B) 1:2 boric acid/compound **8**, C) 1:1 boric acid/compound **8**, D) boric acid buffer standard. All spectra were recorded in 95/5 $H_2O/D_2O$ mixture.

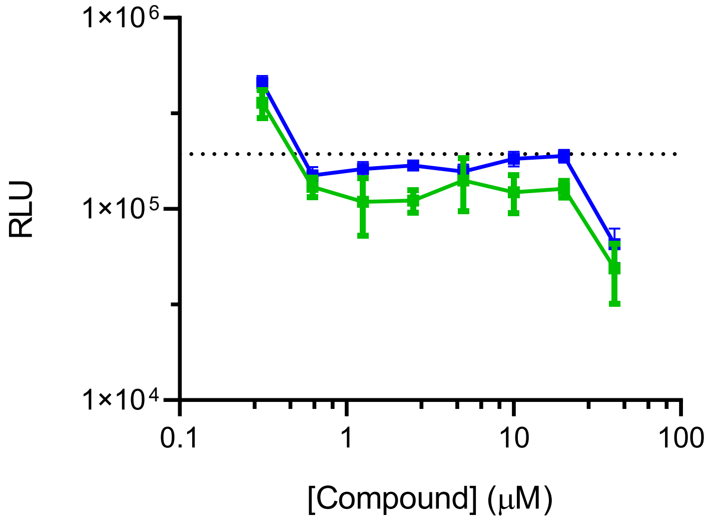


**Figure S15**. Bioluminescence output from the *V. harveyi* TL-26 bioassay strain in response to **10** (blue) and **16** (green). Error bars denote standard deviations for biological replicates, $n$ = 3.

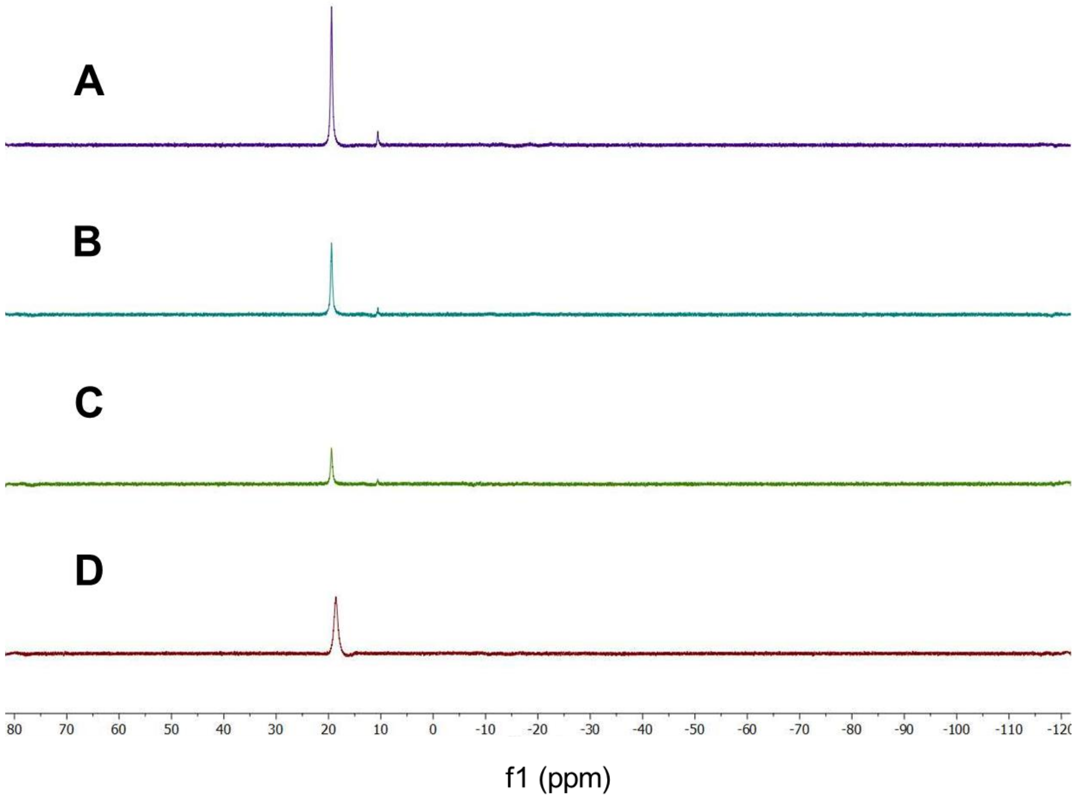


**Figure S16**. $^{11}$B NMR comparison of 12 mM L-xylulose (compound **18**) titrated with different concentrations of boric acid. A) 2:1 boric acid/compound **18**, B) 1:1 boric acid/compound **18**, C) 1:2 boric acid/compound **18**, D) boric acid buffer standard. The peak at 19.4 ppm corresponds to borate. A small peak at 10.6 ppm is apparent when xylulose is present, consistent with the formation of a xylulose borate complex.

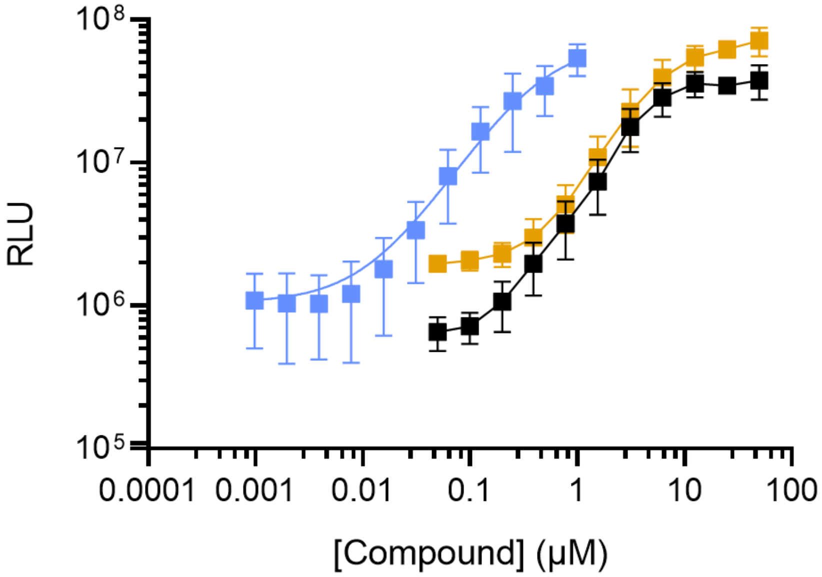


| | EC50 (μM) |
|---|---|
| Compound **1** | 0.4 |
| Compound **8** | 3.4 |
| Compound **18** | 6.0 |

**Figure S17**. Direct comparison of measured EC50 values of AI-2 and associated mimics in *V. harveyi* TL26 bioassay. Light output is shown in response to indicated amounts of compounds **1** (blue), **8** (black), and **18** (orange). Error bars represent standard deviations of technical replicates, n = 3.

**Table S1.** Significant molecular features identified in reactivity-based metabolomics workflow. Observed proton adduct [M+H], retention time (RT), raw abundance, fold change, and qvalue are shown.

| *m/z* | RT (min) | Average Abundance (compound **5** in PBS dataset) | Average Abundance (mimic + compound **5** dataset) | Fold-Change | log2-fold | qvalue |
|---|---|---|---|---|---|---|
| 175.0293 | 6.154642 | 1447769.385 | 16578944.9 | 11.45137 | 3.517448 | 0.005657932 |
| 176.0371 | 5.662175 | 57839.67554 | 1490778.225 | 25.77432 | 4.687862 | 0.00702287 |
| 193.0698 | 6.157067 | 15226.33997 | 157053.8592 | 10.31462 | 3.366618 | 0.005865608 |
| 205.0455 | 4.035233 | 4082.359431 | 841678.6448 | 206.1746 | 7.687722 | 0.009312385 |
| 205.0457 | 5.918442 | 107133.2131 | 9379938.709 | 87.55398 | 6.452101 | 0.008011735 |
| 206.0491 | 5.916925 | 9785.030172 | 975375.0917 | 99.68034 | 6.639237 | 0.007383072 |
| 228.9814 | 5.951092 | 2640.859709 | 275591.4874 | 104.3567 | 6.70538 | 0.001939699 |
| 235.0604 | 5.8959 | 8329.691444 | 700092.3232 | 84.04781 | 6.393138 | 0.004965719 |
| 235.0611 | 5.662258 | 43623.50743 | 3116981.557 | 71.45188 | 6.1589 | 0.009586139 |
| 236.0643 | 5.661342 | 5954.021736 | 384370.5411 | 64.55646 | 6.012489 | 0.009767693 |
| 247.0621 | 5.12125 | 8822.884968 | 548536.3721 | 62.172 | 5.958193 | 0.005239107 |
| 251.0939 | 6.27715 | 11300.7799 | 604859.7812 | 53.52372 | 5.742106 | 0.002042412 |
| 253.1211 | 8.300392 | 163869.2106 | 9951.560014 | 16.46669 | -4.04148 | 0.005222014 |
| 254.0771 | 5.663183 | 11895.98731 | 872676.458 | 73.35889 | 6.1969 | 0.00932076 |
| 265.0749 | 5.5545 | 8156.206224 | 654245.6078 | 80.21445 | 6.32579 | 0.008802364 |
| 265.0749 | 5.1154 | 26201.47268 | 2552119.838 | 97.40368 | 6.605904 | 0.009245141 |
| 268.9739 | 6.957142 | 1206809.672 | 24092.07542 | 50.09156 | -5.6465 | 0.001647105 |
| 283.0874 | 5.55685 | 16121.94232 | 3484077.422 | 216.1078 | 7.755607 | 0.008927702 |
| 283.1247 | 5.110192 | 32363.98125 | 800851.5304 | 24.74515 | 4.629074 | 0.001737254 |
| 296.0601 | 5.946483 | 2714.484245 | 391444.3905 | 144.2058 | 7.171985 | 0.006714692 |
| 297.104 | 5.776733 | 2287.018179 | 1311307.642 | 573.37 | 9.163323 | 0.006770262 |
| 298.107 | 5.747967 | 752.4114117 | 196617.0145 | 261.3158 | 8.029651 | 0.008540012 |
| 350.9769 | 6.979908 | 500007.7689 | 1566.322303 | 319.2241 | -8.31843 | 0.006201043 |
| 355.1315 | 5.884517 | 1735.116785 | 4851601.044 | 2796.124 | 11.44921 | 0.004105654 |
| 363.0576 | 5.14435 | 1553.706522 | 1603381.713 | 1031.972 | 10.01119 | 0.009534706 |
| 393.0692 | 4.89445 | 274.8622079 | 4751215.462 | 17285.81 | 14.0773 | 0.004884196 |
| 434.0956 | 5.304133 | 186.3713771 | 1232156.327 | 6611.296 | 12.69072 | 0.004034585 |

**Table S2.** Top 10 most abundant fragment ions in MSMS compound **5**-derivatized AI-2 mimic. Collision energy = 20 eV.

| Height | *m/z* |
|---|---|
| 11969.37 | 175.0295 |
| 14399.21 | 189.0483 |
| 16267.88 | 206.0499 |
| 21506.45 | 219.0644 |
| 23697.15 | 247.0629 |
| 24766.35 | 203.0304 |
| 25215.67 | 201.0506 |
| 34137.32 | 265.0756 |
| 34171.21 | 176.0375 |
| 36435.18 | 235.0616 |
| 90029.63 | 205.0464 |

**Table S3.** Top 10 most abundant fragment ions in MSMS compound **5**-derivatized synthetic L-xylosone (compound **8**). Collision energy = 20 eV.

| Height | *m/z* |
|---|---|
| 12426.39 | 189.0487 |
| 16464.8 | 175.0299 |
| 21282.8 | 206.05 |
| 24707.37 | 219.0644 |
| 25019.65 | 247.0635 |
| 25350.53 | 201.0506 |
| 29678.37 | 176.0379 |
| 29910.82 | 203.0304 |
| 35991.71 | 265.0761 |
| 41043.22 | 235.0619 |
| 145570.7 | 205.0465 |

*Synthesis of compounds **8** and **9***

*Step 1: Synthesis of Compound **13***

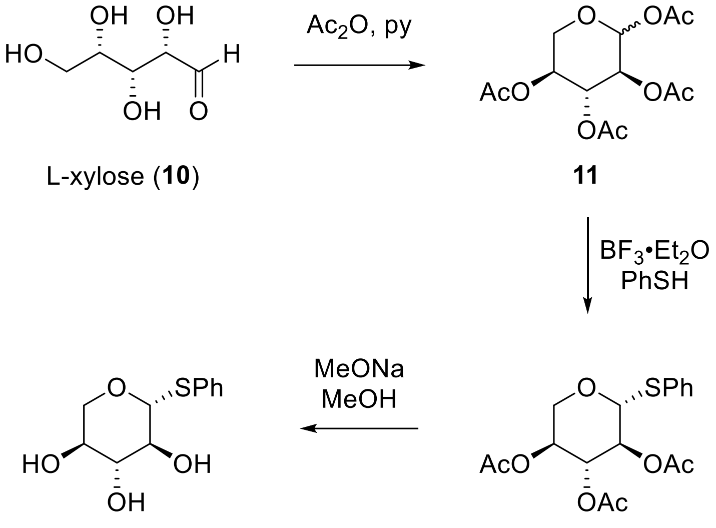


To a 250 mL round-bottom flask containing compound **10** (20 mmol, 3.00 g) was added Ac2O (200 mmol, 18.9 mL, 10 equiv) and pyridine (200 mmol, 10 equiv, 16.1 mL). The mixture was heated at 80° C for 2 h. After cooling to rt, MeOH (10 mL) was added and stirred for another 30 min. The mixture was diluted in EtOAc (200 mL), washed with 1 M HCl (200 mL), $H_2O$ (200 mL), satd. $NaHCO_3$ (200 mL) and brine (200 mL), dried over $Na_2SO_4$, filtered and concentrated *in vacuo* to afford compound **11**.

To a solution of compound **11** prepared above in dry $CH_2Cl_2$ (30 mL) were added thiophenol (2.46 mL, 24 mmol, 1.2 equiv) and $BF_3{\cdot}Et_2O$ (7.37 mL, 60 mmol, 3.0 equiv) at 0 °C under argon atmosphere. The solution was stirred for 2 h at room temperature and then diluted with $CH_2Cl_2$ (20 mL). The mixture was neutralized by the addition of satd. $NaHCO_3$ (100 mL), extracted with $CH_2Cl_2$ (2×50 mL), dried over $Na_2SO_4$, filtered and concentrated *in vacuo*. The residue was dissolved in MeOH (20 mL) and MeONa (4 mmol, 1.0 M in MeOH, 4 mL) was added. After stirring for 15 min at rt, the mixture was neutralized with $NH_4Cl$ (5 mmol, 267.5 mg) and concentrated *in vacuo*. The dried material was redissolved in $Me_2CO$, filtered, and concentrated *in vacuo*. After recrystallization of the residue from acetone-hexanes, compound **13** was obtained (3.49 g, 72% over 3 steps) as an off-white solid.

$^1$H-NMR (500 MHz, acetone-d$_6$) δ 7.50 (d, *J* = 7.6 Hz, 2H), 7.32 (t, *J* = 7.5 Hz, 2H), 7.26 (t, *J* = 7.4 Hz, 1H), 4.71 (d, *J* = 8.5 Hz, 1H), 4.38 (dd, *J* = 11.9, 4.5 Hz, 1H), 4.22 (d, *J* = 4.5 Hz, 1H), 3.98 (dd, *J* = 11.3, 4.7 Hz, 1H), 3.52 (td, *J* = 8.7, 4.5 Hz, 1H), 3.49 – 3.41 (m, 1H), 3.35 – 3.26 (m, 2H). The characterization data we obtained match those reported previously.[1]

*Step 2: Acetonide protection of Compound **13***

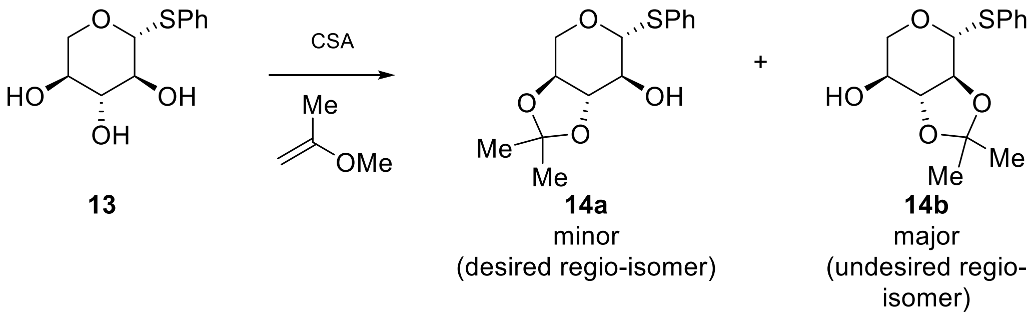


To a 100 mL round bottom flask containing the triol compound **13** (2.04 g, 8.26 mmol) and (+)-camphor sulphonic acid (CSA, 192.0 mg, 0.82 mmol) in dry DMF (40 mL), 2-methoxypropene (2.38 mL, 24.8 mmol) was added dropwise under nitrogen atmosphere. The resulting mixture was stirred at rt for 2 h. Saturated sodium bicarbonate solution (160 mL) was added to the reaction mixture and stirred for another 30 min. The mixture was diluted with EtOAc (300 mL), washed with water (150 mL), brine (150 mL), dried over $Na_2SO_4$ and concentrated *in vacuo*. The crude product was purified on $SiO_2$ (Hexanes/EtOAc 10:1 to 3:1) to afford both regio-isomers of the acetonide protected sugar (major isomer: 1.78 g, 6.31 mmol, 76%, off-white solid; minor isomer: 223.2 mg 0.79 mmol, 10%, off-white solid).

Compound **14b** (major regio-isomer): $^1$H-NMR (500 MHz, acetone-$d_6$) δ 7.53 (d, *J* = 6.5 Hz, 2H), 7.42 – 7.25 (m, 3H), 4.94 (d, *J* = 9.5 Hz, 1H), 4.57 (d, *J* = 4.8 Hz, 1H), 3.98 (dd, *J* = 11.5, 5.3 Hz, 1H), 3.91 – 3.79 (m, 1H), 3.59 (t, *J* = 9.1 Hz, 1H), 3.26 (dd, *J* = 11.5, 9.1 Hz, 1H), 3.18 (t, *J* = 9.1 Hz, 1H), 1.39 (s, 3H), 1.36 (s, 3H); $^{13}$C-NMR (126 MHz, acetone-$d_6$) δ 133.8, 132.8, 129.7, 128.4, 110.8, 85.5, 83.7, 76.3, 71.0, 69.6, 27.0, 26.7. The characterization data we obtained match those reported previously.[2]

Compound **14a** (minor regio-isomer**):** $^1$H-NMR (500 MHz, acetone-$d_6$) δ 7.54 (d, *J* = 7.4 Hz, 2H), 7.33 (t, *J* = 7.3 Hz, 2H), 7.29 (t, *J* = 7.2 Hz, 1H), 4.75 (d, *J* = 4.4 Hz, 1H), 4.59 (d, *J* = 8.5 Hz, 1H), 4.13 (dd, *J* = 9.8, 4.4 Hz, 1H), 3.65 – 3.56 (m, 2H), 3.54 (t, *J* = 9.1 Hz, 1H), 3.48 – 3.40 (m, 1H), 1.37 (s, 6H); $^{13}$C-NMR (126 MHz, acetone-$d_6$) δ 134.9, 132.3, 129.7, 128.1, 111.4, 90.0, 83. 9, 74.2, 72.7, 68.5, 27.0, 26.8.

*Step 3: Synthesis of Compound **15***

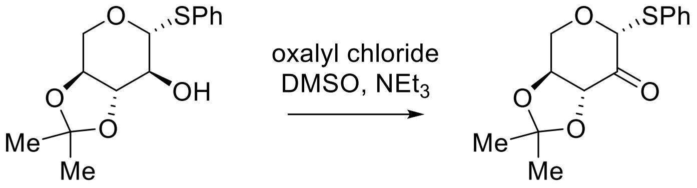


**14a** **15**

Oxalyl chloride (70 µL, 0.8 mmol) was dissolved in $CH_2Cl_2$ (4 mL) in a flame-dried 25 mL round-bottom flask and cooled to -78 °C. Dimethyl sulfoxide (DMSO, 70 µL, 1.0 mmol) was added dropwise and allowed to stir for 15 min. A solution of compound **14a** (112.9 mg, 0.4 mmol) in $CH_2Cl_2$ (2 mL) was added and stirred at -78 °C for 1 h followed by $NEt_3$ (167 µL, 1.2 mmol). The reaction mixture was stirred for another 3 h at -78°C and quenched by the addition of $H_2O$ (10 mL). The aqueous layer was extracted with $CH_2Cl_2$ (3 x 10 mL), washed with brine (10 mL), dried over $Na_2SO_4$ and concentrated *in vacuo*. The crude product was purified on $SiO_2$ (Hexanes/EtOAc 10:1 to 5:1) to afford the ketone product **15** (71.8 mg, 0.26 mmol, 64% yield) as an off-white solid.

$^1$H-NMR (500 MHz, acetone-$d_6$) δ 7.57 (d, $J$ = 6.6 Hz, 2H), 7.44 – 7.37 (m, 3H), 5.73 (s, 1H), 4.71 (d, $J$ = 6.1 Hz, 1H), 4.66 – 4.57 (m, 2H), 4.05 (d, $J$ = 13.7 Hz, 1H), 1.38 (s, 3H), 1.36 (s, 3H); $^{13}$C-NMR (126 MHz, acetone-$d_6$) δ 200.2, 137.2, 133.3, 130.2, 129.1, 111.2, 90.2, 77.2, 76.2, 61.7, 27.3, 26.0.

*Step 4: Synthesis of L-xylosone (**8**)*

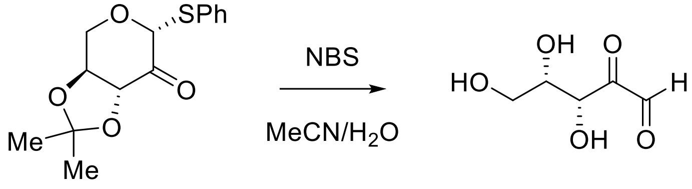


**15** L-xylosone (**8**)

The protected ketone **15** (42.1 mg, 0.15 mmol) was dissolved in MeCN (1 mL) and $H_2O$ (1 mL). N-Bromosuccinimide (NBS, 30.0 mg, 0.165 mmol) was added. The resulting mixture was stirred at rt for 10 min and quenched by the addition of NaOAc (1 mL, 1 M in $H_2O$). The mixture was concentrated *in vacuo* and purified on C18 modified $SiO_2$ ($H_2O$ to 10% MeCN in $H_2O$).

The $^1$H-NMR spectrum of xylosone appears to be complicated due to complex speciation of the sugar in water. The identity and purity of xylosone was determined by derivatization with diphenylhydrazine (DPH) shown below. DPH was chosen for this analysis because it results in a more stable product than does derivatizing with compound **5** used in the metabolomics analyses. Thus, derivatization with DPH enabled isolation and purification.

*D-xylosone was prepared and analyzed using the identical sequence described above for synthesis of L-xylosone starting with D-xylose.*

*Diphenylhydrazine (DPH) derivatization of compound **8***

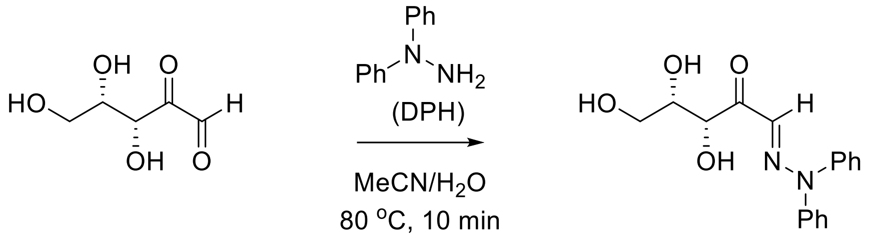


L-xylosone (**8**)

L-xylosone DPH derivative (**19**)

To 200 µL of the resulting fraction from the C18 purification in step 4 was added 200 µL of 0.1 M diphenyl hydrazine (DPH) solution (prepared by dissolving 1 mmol of diphenylhydrazine hydrochloride and 3 mmol of $NEt_3$ in 5 mL $H_2O$ and 5 mL MeCN). The resulting mixture was reacted at rt for 24 h. The resulting mixture was purified on using HPLC to afford the DPH derivatized sugar (**19**).

**$^1$H-NMR** (500 MHz, MeOH-$d_4$) δ 7.49 (brs, 4H), 7.33 (brs, 2H), 7.22 (dd, *J* = 8.6, 1.2 Hz, 4H), 6.62 (s, 1H), 5.26 (d, *J* = 2.1 Hz, 1H), 4.29 (td, *J* = 6.7, 2.1 Hz, 1H), 3.81 (dd, *J* = 10.9, 7.0 Hz, 1H), 3.67 (dd, *J* = 10.9, 6.4 Hz, 1H);

**$^{13}$C-NMR** (126 MHz, MeOH-$d_4$) δ 199.2, 131.2, 75.1, 74.4, 64.0. Aromatic signals are missing from the $^{13}$C-NMR spectrum due to hindered rotation.

**ESI-HRMS** cald. for $C_{17}H_{19}N_2O_4^+$ $[M+H]^+$ = 315.1339 m/z, found 315.1337 m/z. The characterization data we obtained match the ones reported previously.[3]

HPLC method: Kinetex 5 µm C18 100 Å, LC Column 250 x 10.0 mm (70% MeCN in $H_2O$, 2.5 mL/min, 15 min). DPH modified xylosone was eluted between 4.5 and 4.9 min.

*Quantitation of xylosone stock solution*

**<u>Procedure</u>:** The extinction coefficient (ε) of compound **19** was determined using HPLC with UV detection at 254 nm. A 5 mg/mL stock solution was prepared by dissolving purified compound **19** in MeCN. Serial dilutions in MeCN were made to obtain stock solutions with concentrations of 2.5, 1.25, 0.625, and 0.3125 mg/mL. Each solution was filtered through a 0.22 μm membrane before injection.

Purified compound **8** or **9** was dissolved in 10 mL water. A 400 μL aliquot of the solution was derivatized with 200 μL of 0.1 M DPH (prepared by dissolving 1 mmol of diphenylhydrazine hydrochloride and 3 mmol of $NEt_3$ in 5 mL $H_2O$ and 5 mL MeCN), diluted to 1 mL using MeOH and allowed to stand for 24 h at rt before analyzing by HPLC. The concentrations of the initial xylosone solutions were calculated using the standard curve.

HPLC method: Kinetex 5 μm C18 100 Å, LC Column 250 x 10.0 mm (10 μL per injection, 70% MeCN in $H_2O$, 2.5 mL/min, 15 min).

$^1$H-NMR, 500 MHz, acetone-$d_6$

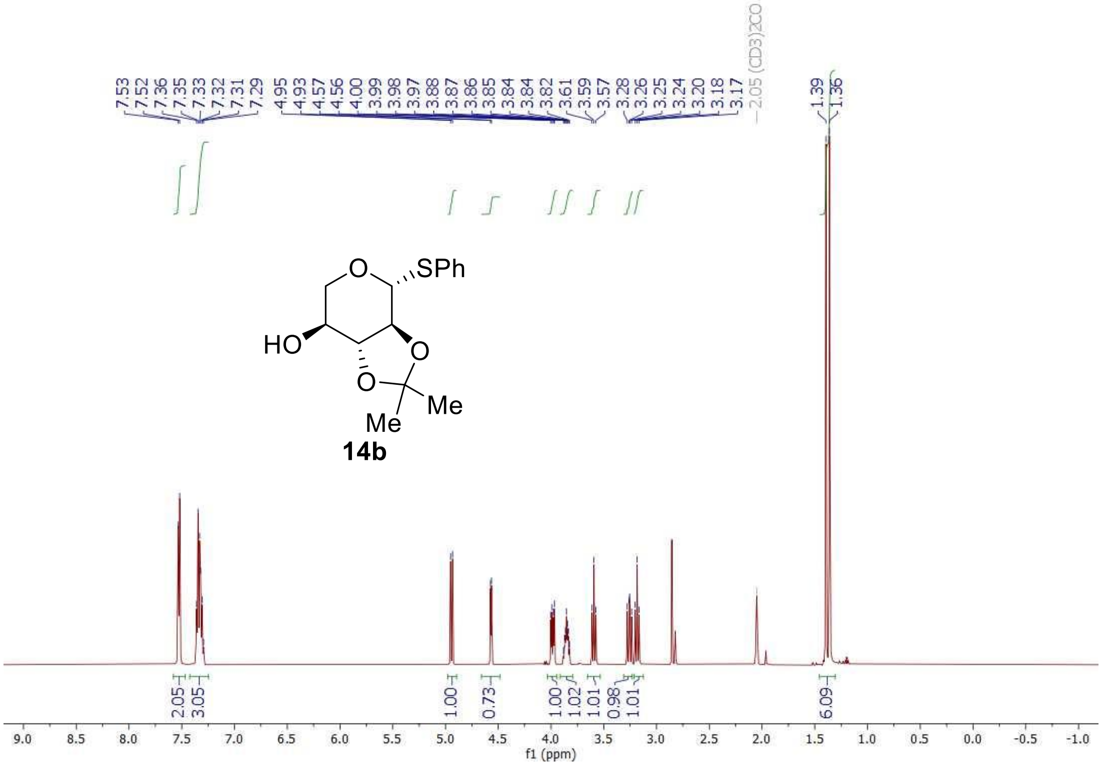

$^{13}$C-NMR, 126 MHz, acetone-d$_6$

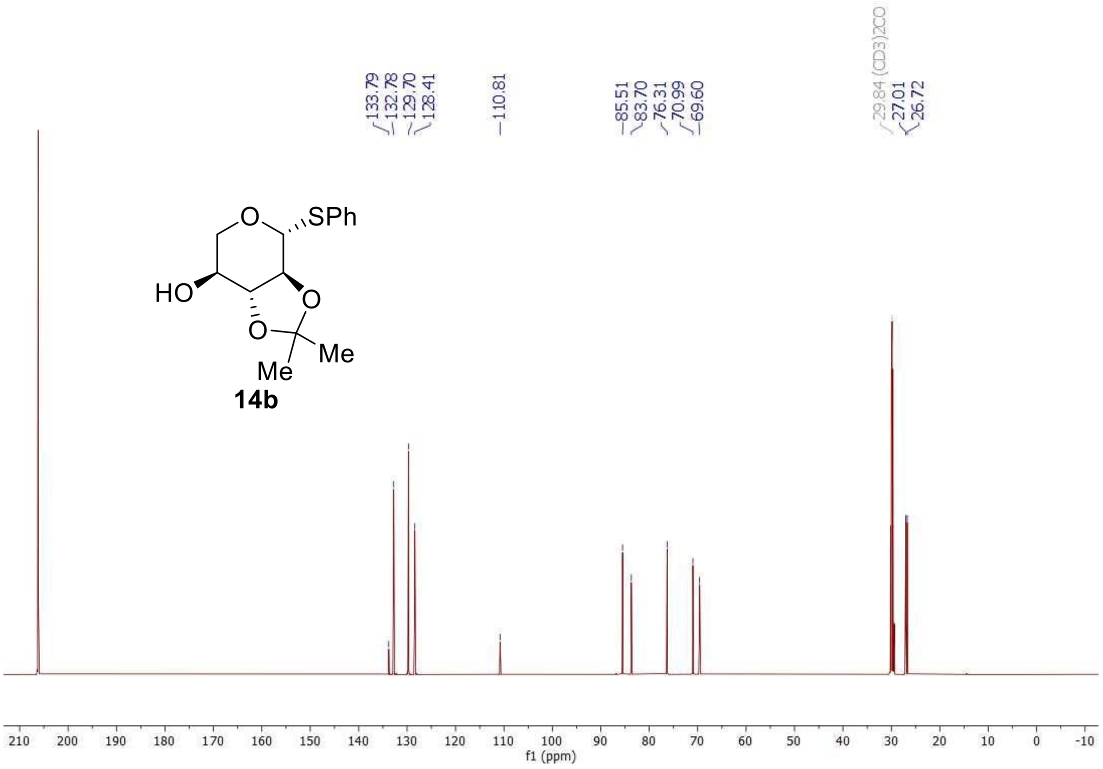

$^1H$-$^1H$ COSY, 500 MHz, acetone-$d_6$

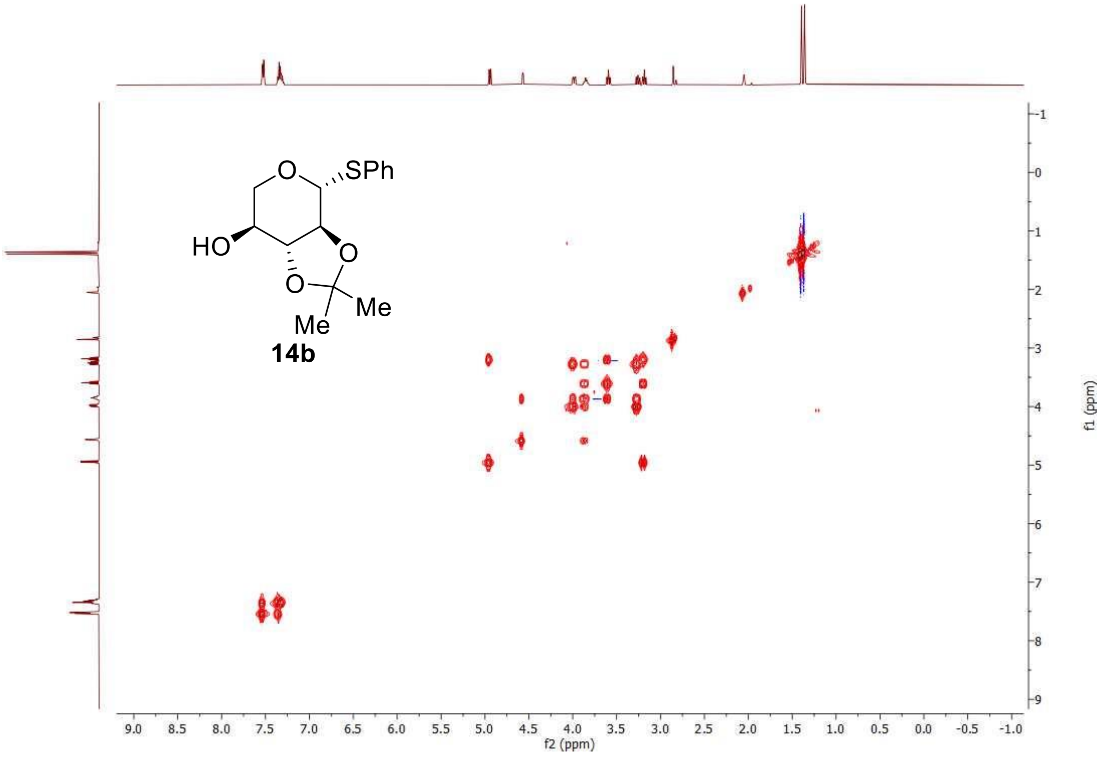

$^{1}H$-$^{13}C$ HSQC, 500 MHz, acetone-$d_6$

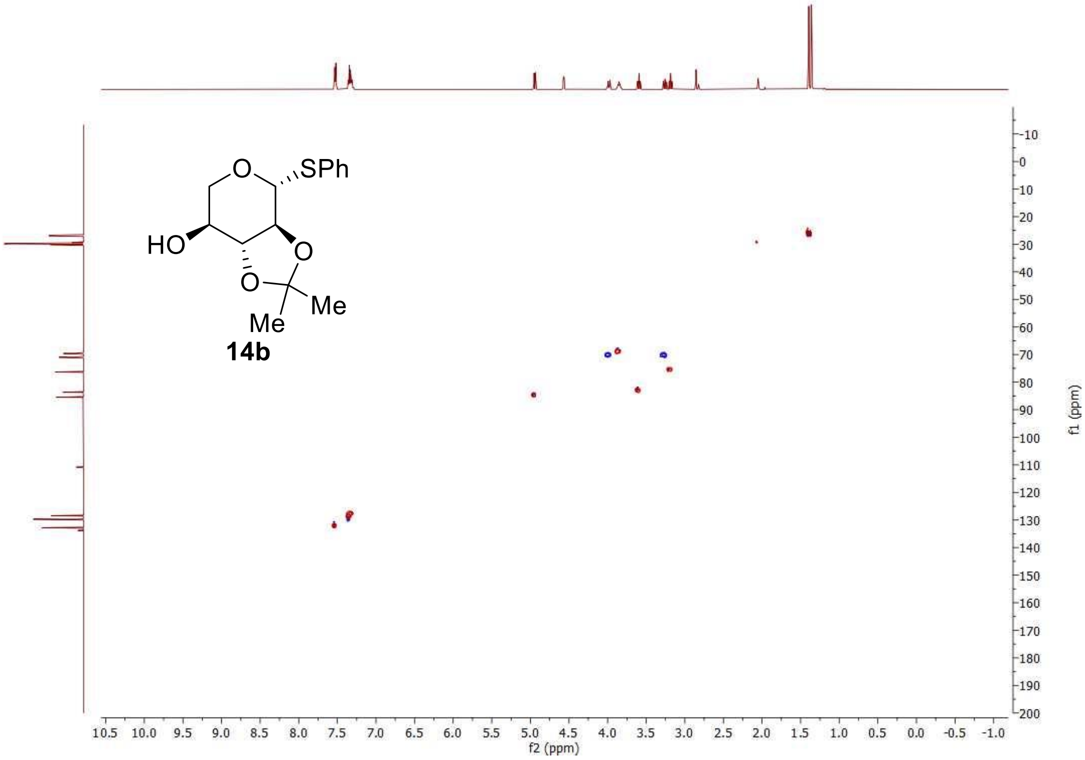

$^{1}H$-$^{13}C$ HMBC, 500 MHz, acetone-$d_6$

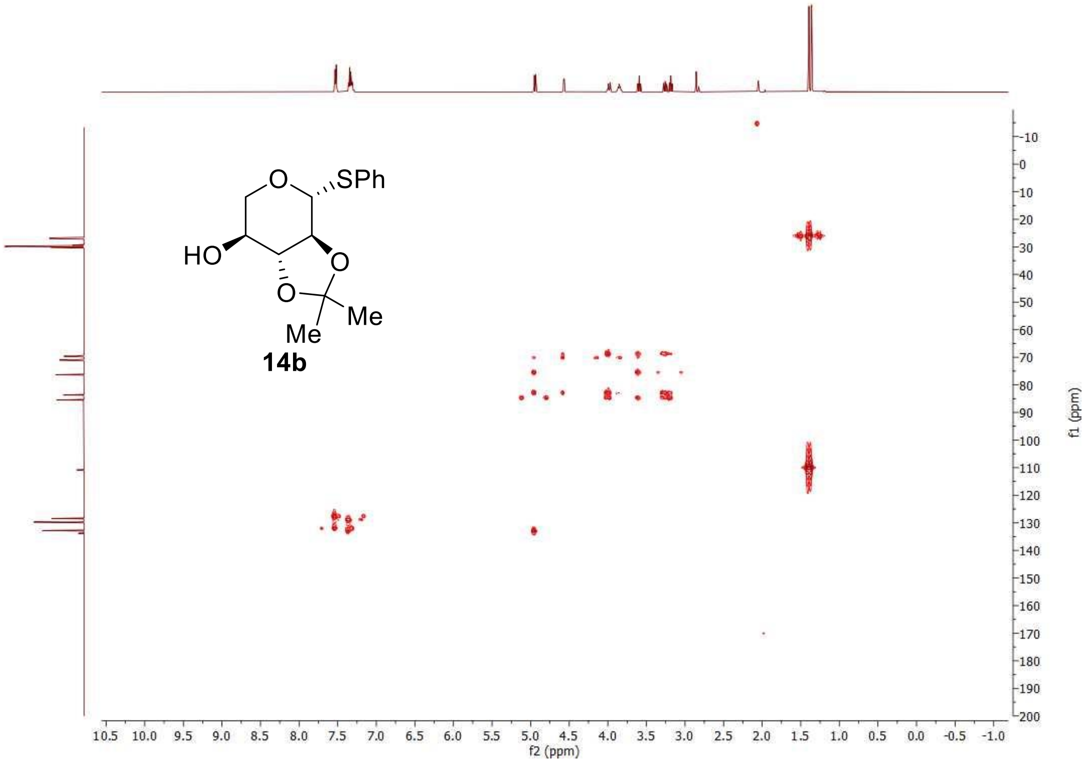

$^{1}$H-NMR, 500 MHz, acetone-d$_6$

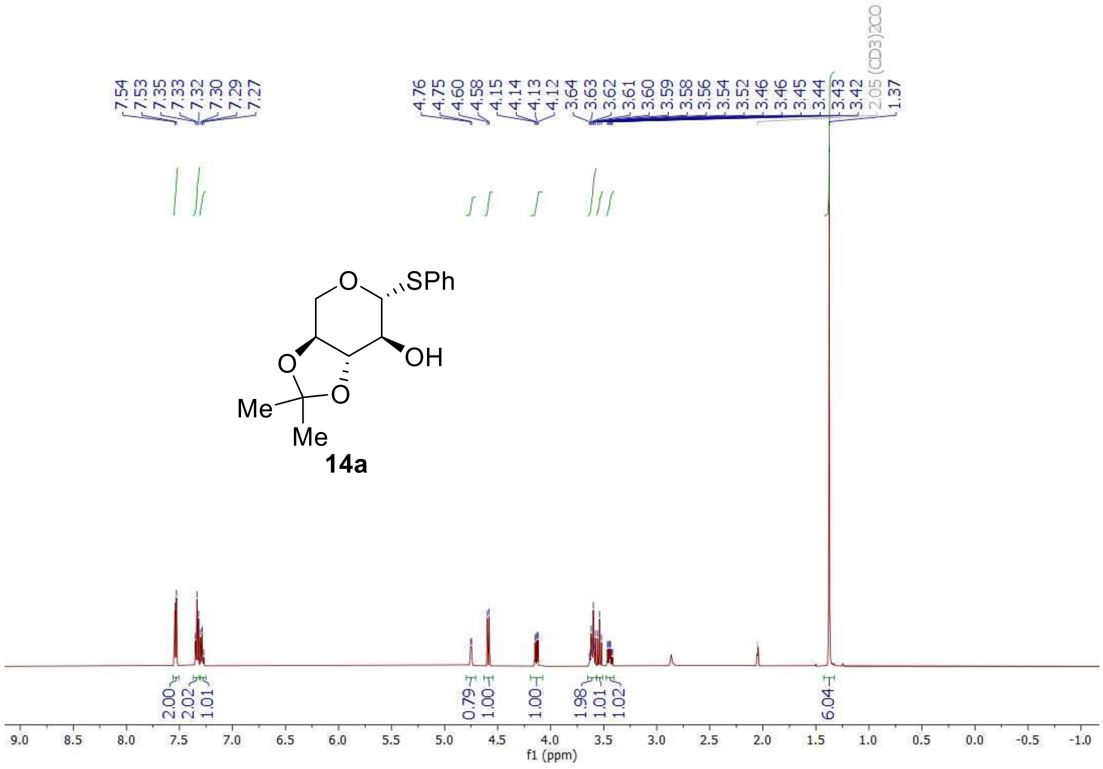

$^{13}$C-NMR, 126 MHz, acetone-$d_6$

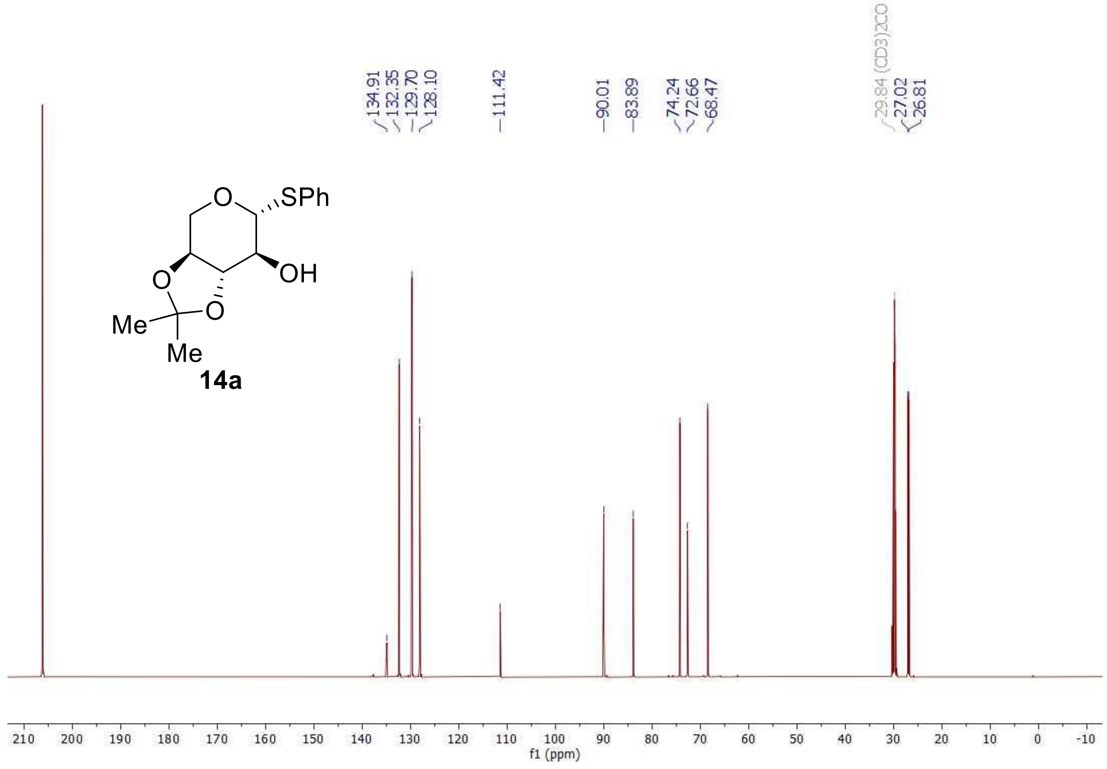

$^{1}H$-$^{1}H$ COSY, 500 MHz, acetone-$d_6$

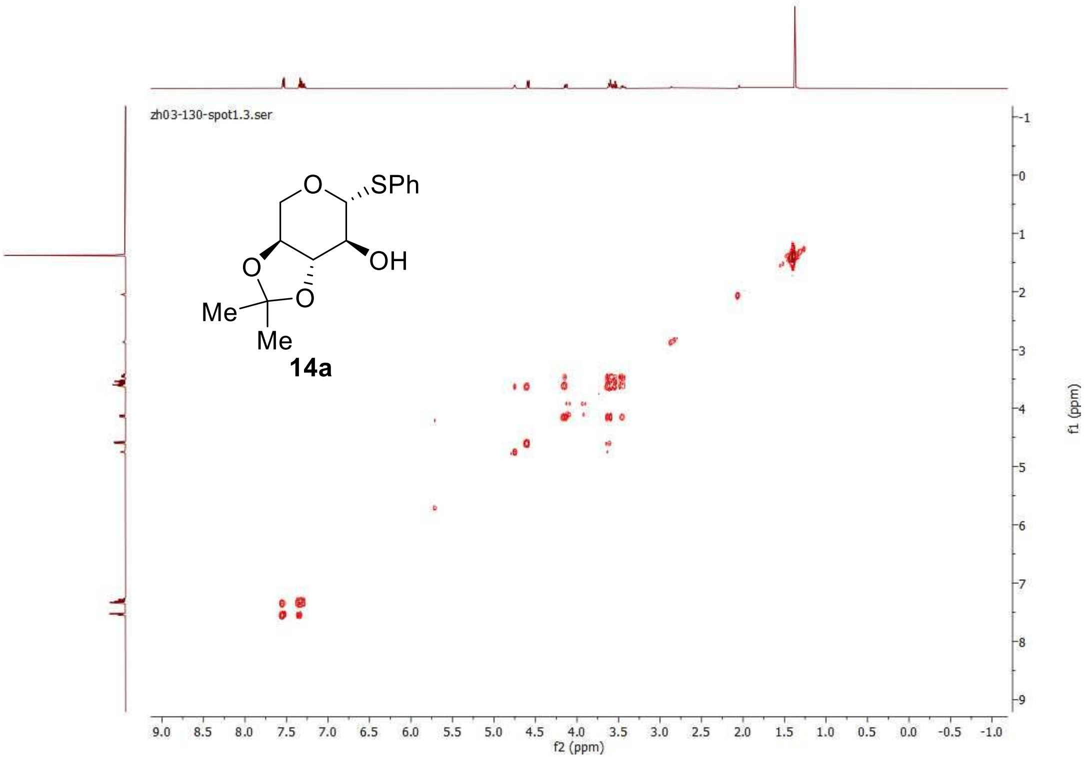

$^{1}H$-$^{13}C$ HSQC, 500 MHz, acetone-$d_6$

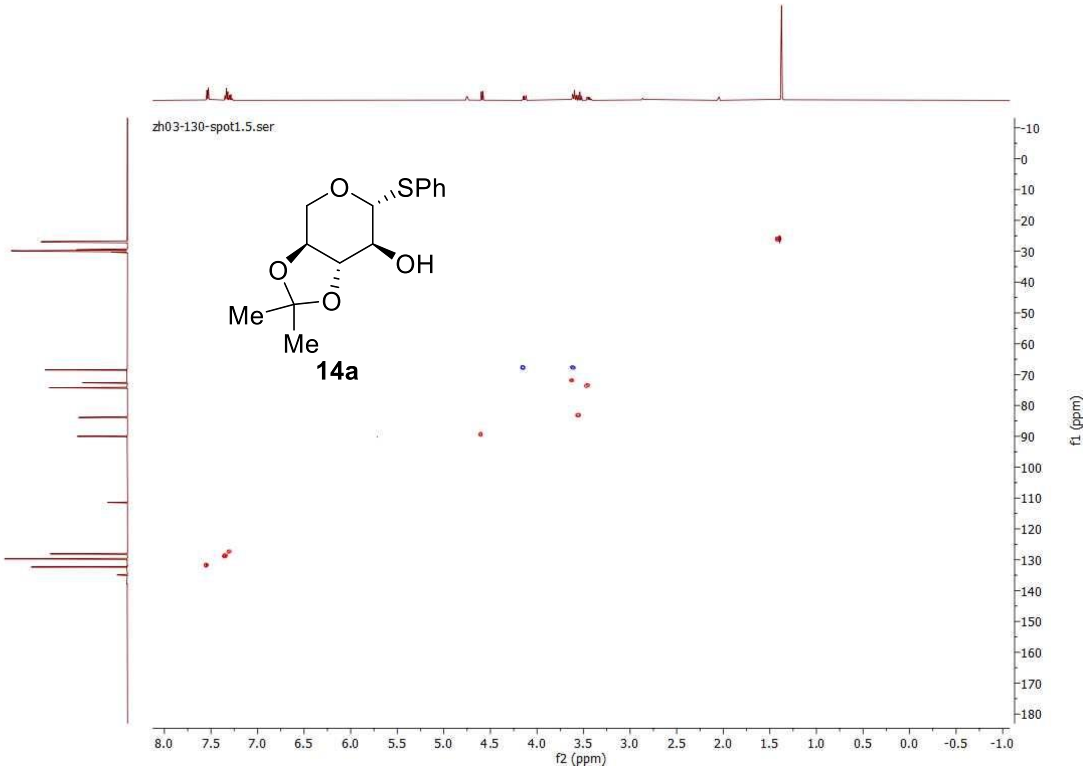

$^{1}H$-$^{13}C$ HMBC, 500 MHz, acetone-$d_6$

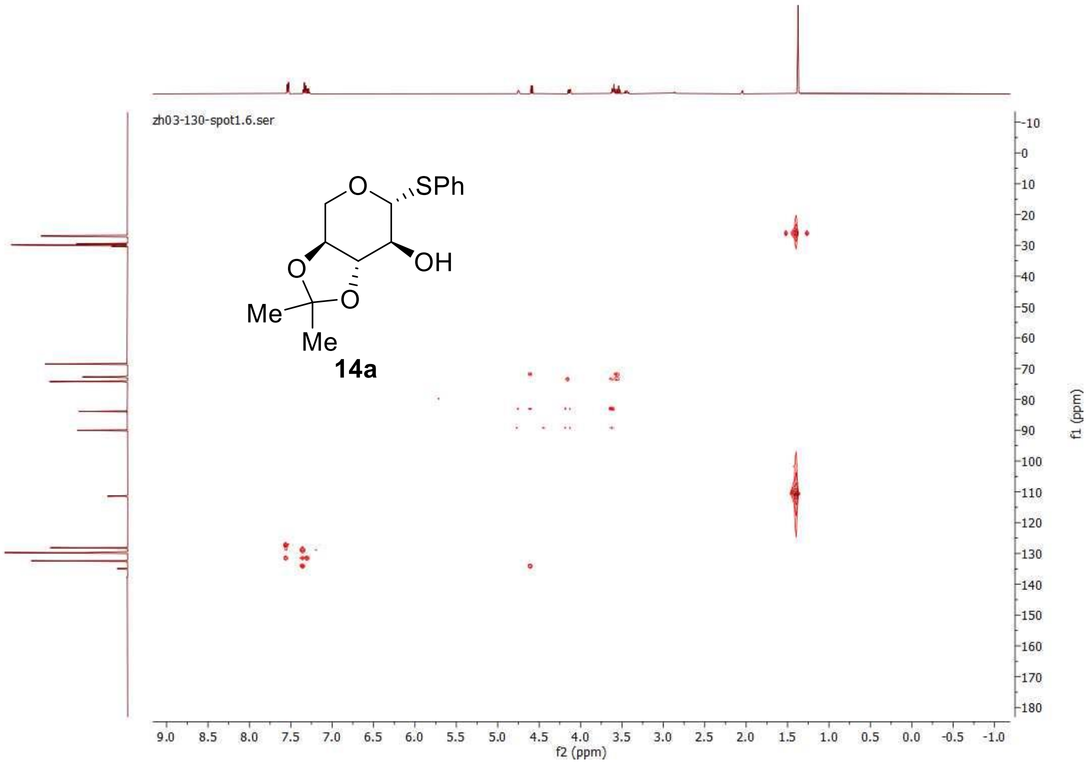

$^{1}$H-NMR, 500 MHz, acetone-$d_6$

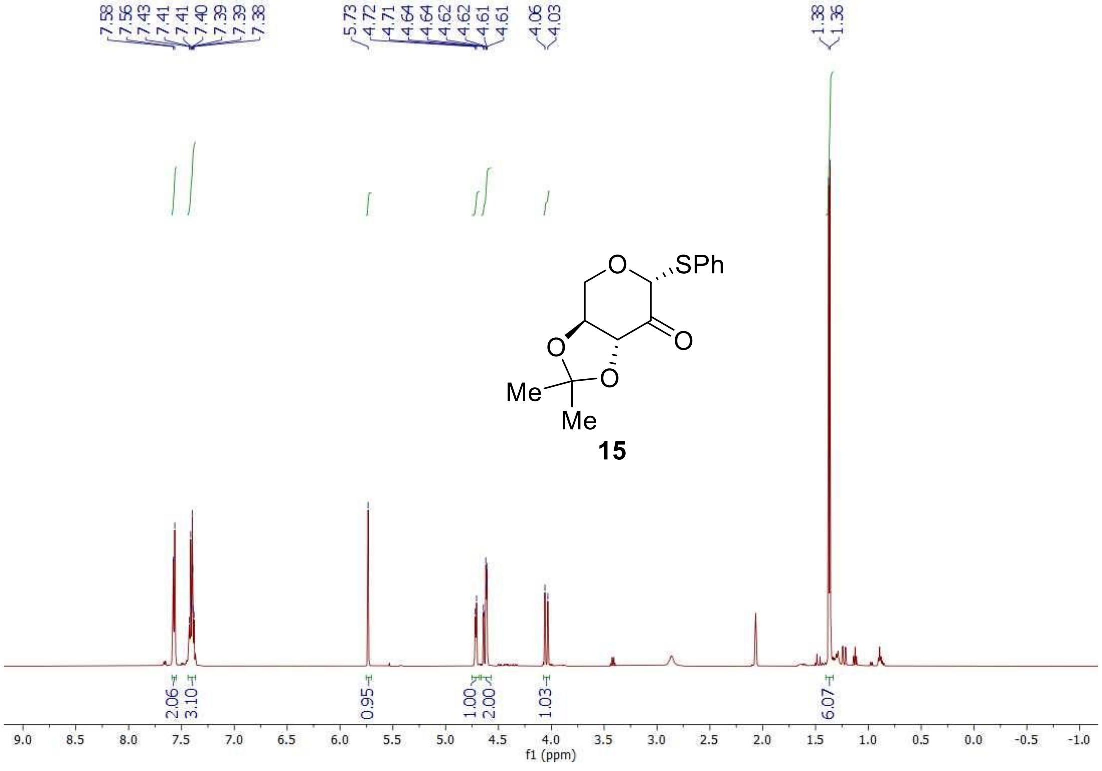

<sup></sup>$^{13}$C-NMR, 126 MHz, acetone-$d_6$

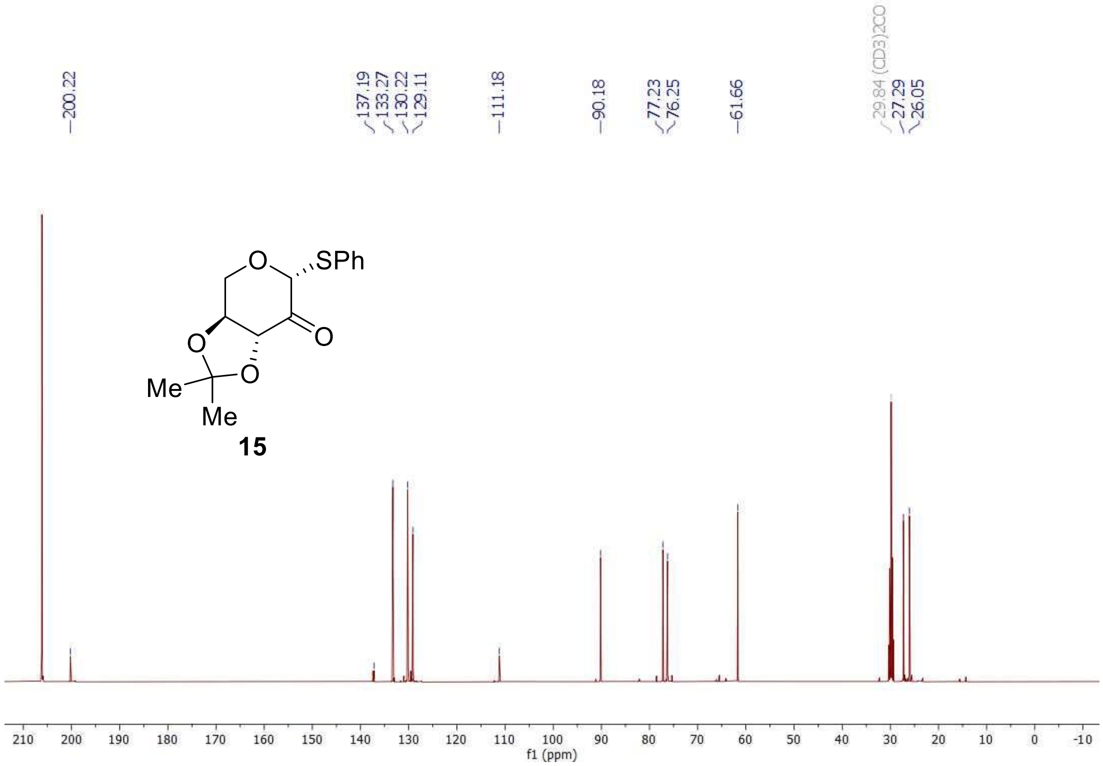

$^{1}H$-NMR, 500 MHz, $D_2O$

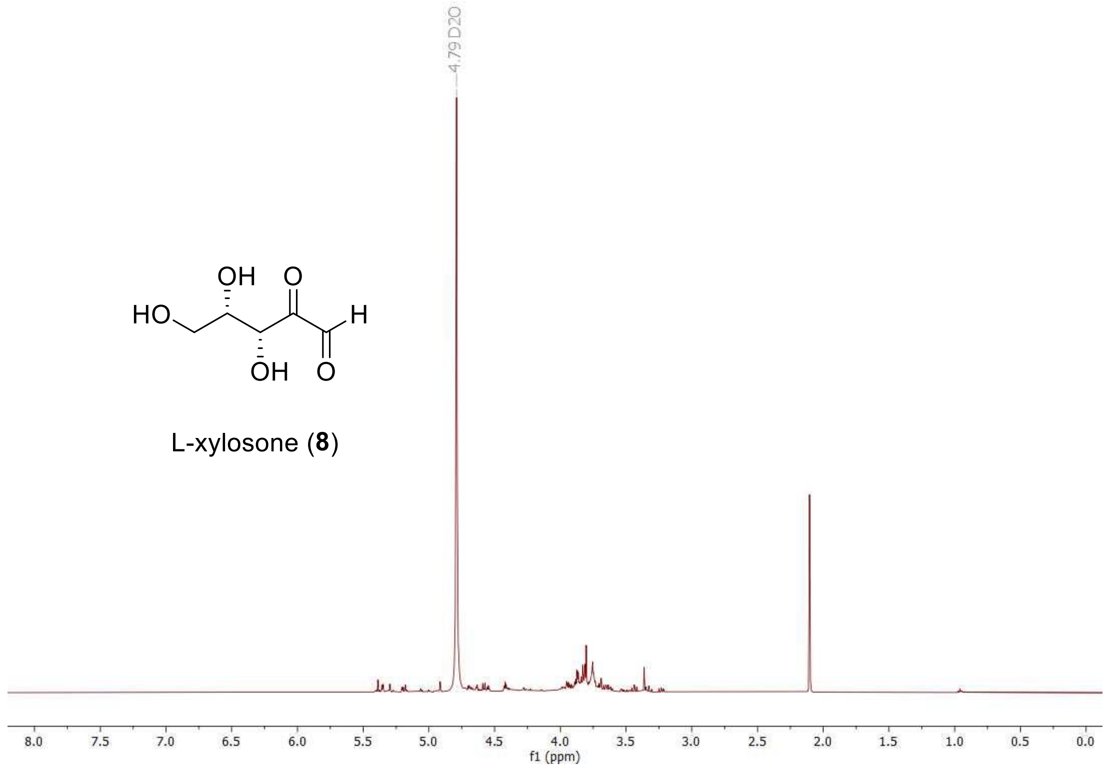

[1]H-NMR, 500 MHz, MeOH-d4

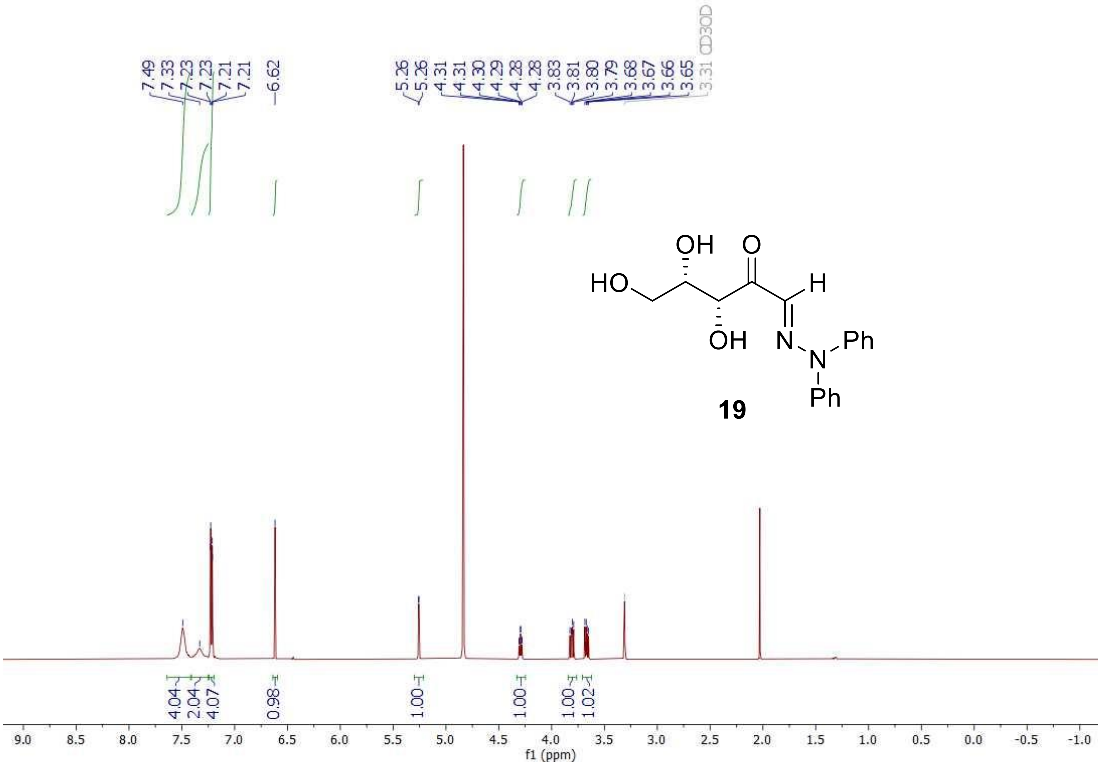

$^{13}$C-NMR, 126 MHz, MeOH-d$_4$

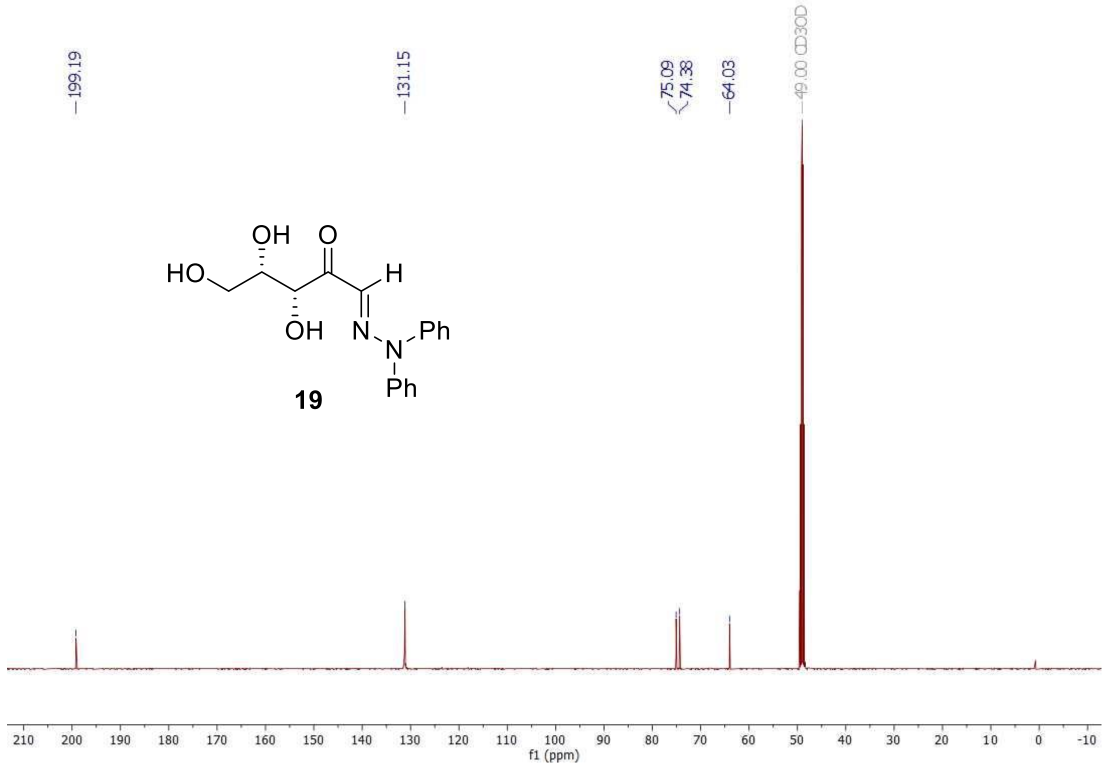

$^{1}H$-$^{1}H$ COSY, 500 MHz, MeOH-$d_4$

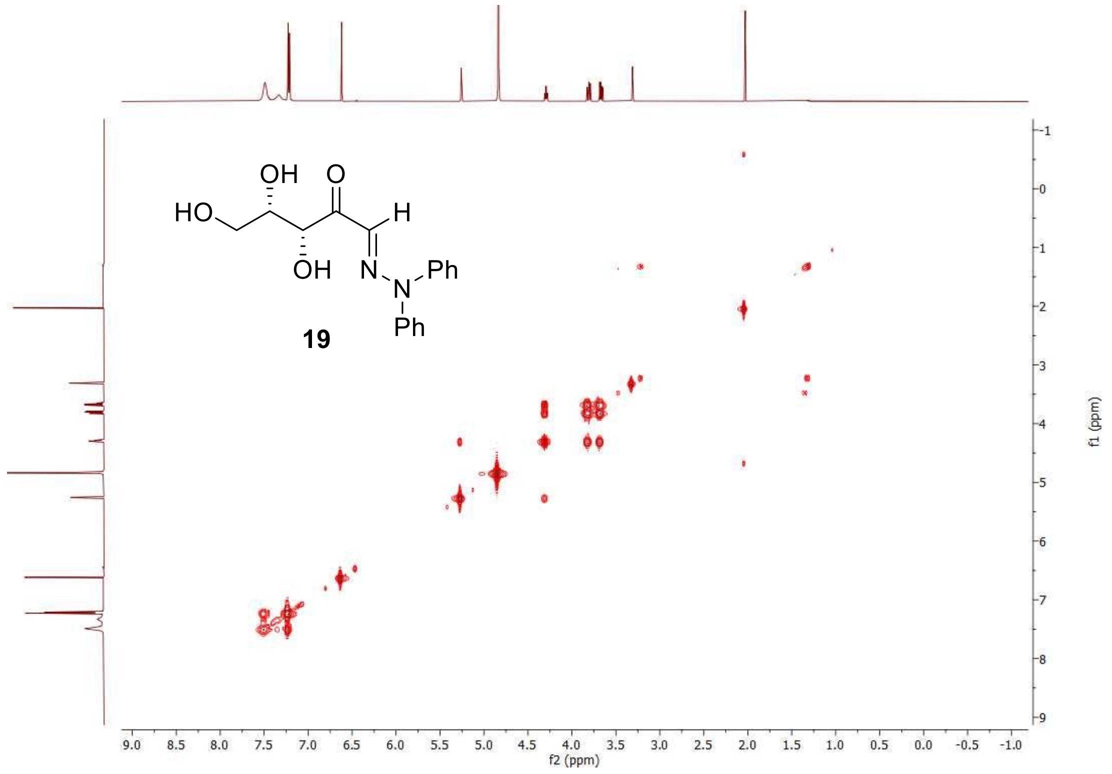

$^{1}H$-$^{13}C$ HSQC, 500 MHz, MeOH-$d_4$

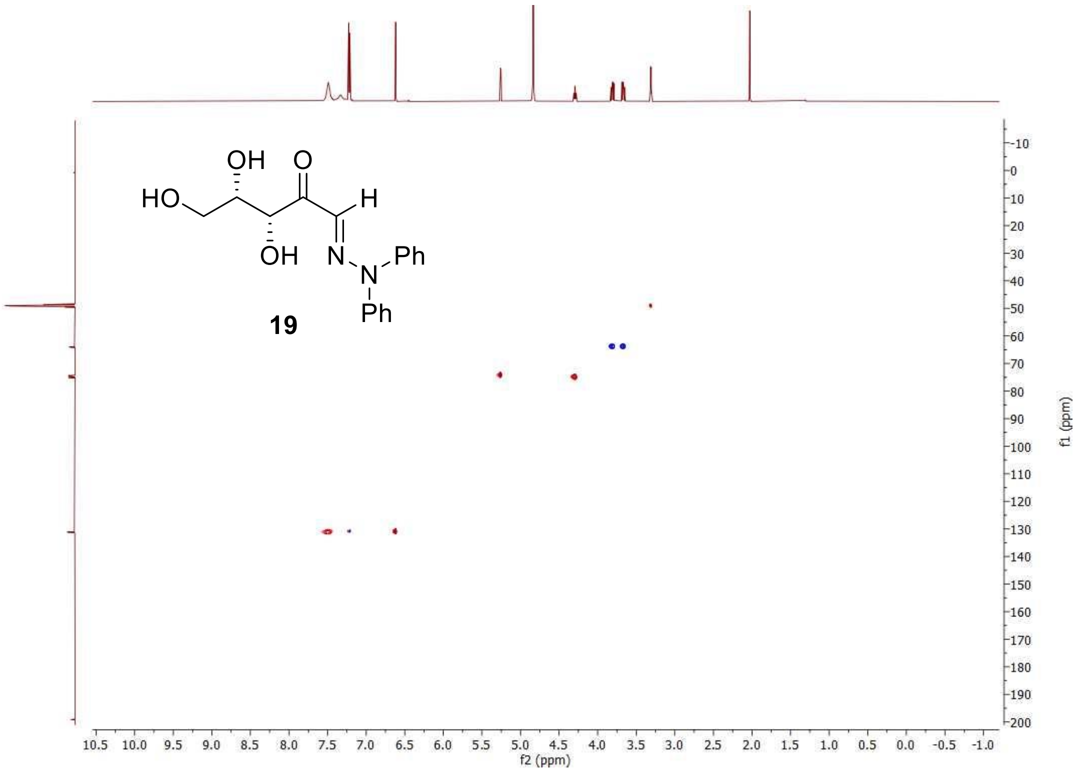

$^{1}H$-$^{13}C$ HMBC, 500 MHz, MeOH-$d_4$

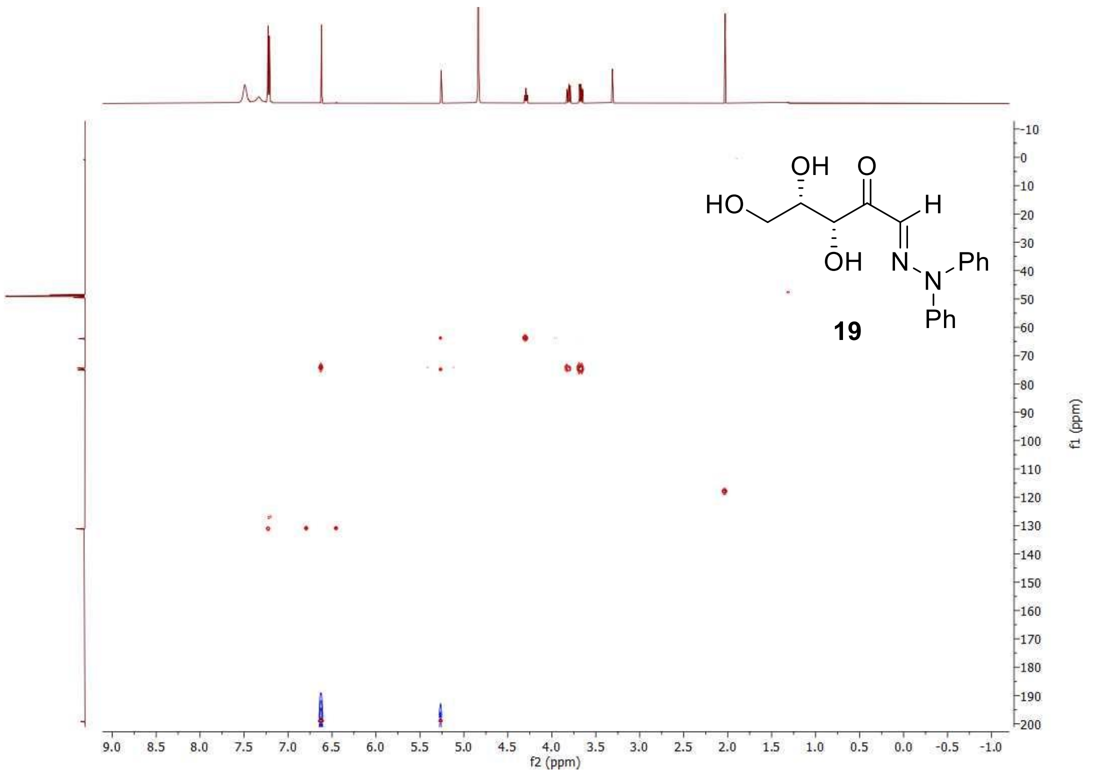

### Supplementary References


1. Lopez, R. & Fernandez-Mayoralas, A. Enzymic .beta.-Galactosidation of Modified Monosaccharides: Study of the Enzyme Selectivity for the Acceptor and Its Application to the Synthesis of Disaccharides. *J. Org. Chem.* **59**, 737-745 (1994).
2. Phanumartwiwath, A., Hornsby, T. W., Jamalis, J., Bailey, C. D. & Willis, C. L. Silyl Migrations in d-Xylose Derivatives: Total Synthesis of a Marine Quinoline Alkaloid. *Org. Lett.* **15**, 5734-5737 (2013).
3. Volc, J., Sedmera, P., Halada, P., Přikrylová, V. & Haltrich, D. Double oxidation of d-xylose to d-glycero-pentos-2,3-diulose (2,3-diketo-d-xylose) by pyranose dehydrogenase from the mushroom Agaricusbisporus. *Carbohydr. Res.* **329**, 219-225 (2000).
